# Impact of glycan positioning on HIV-1 Env glycan shield density, function, and antibody recognition

**DOI:** 10.1101/2020.03.31.019091

**Authors:** Qing Wei, Audra A. Hargett, Barbora Knoppova, Alexandra Duverger, Reda Rawi, Chen-Hsiang Shen, S. Katie Farney, Stacy Hall, Rhubell Brown, Brandon F. Keele, Sonya L. Heath, Michael S. Saag, Olaf Kutsch, Gwo-Yu Chuang, Peter D. Kwong, Zina Moldoveanu, Milan Raska, Matthew B. Renfrow, Jan Novak

## Abstract

N-glycans, which represent >50% mass of the HIV-1 envelope (Env) trimer, play important roles for virus-cell entry and immune evasion. How each glycan unit interacts to shape the Env protein-sugar complex and affects Env function is not well understood. Here, high-resolution glycomics analysis of two Env variants from the same donor, with differing functional characteristics and N-glycosylation-site composition, revealed that changes to key N-glycosylation-site not only affected the Env structure at distant locations, but also had a ripple effect on Env-wide glycan processing, virus infectivity, and antibody recognition and virus neutralization. Specifically, the N262 glycan, although not located in the CD4-binding site, controlled Env binding to the CD4 receptor, affected the recognition of Env by several glycan-dependent broadly neutralizing antibodies, and altered heterogeneity of glycosylation at several sites, with N156, N160, and N448 displaying limited glycan processing. Molecular dynamic simulations visualized how specific oligosaccharide positions can move to compensate for loss of a glycan. This study demonstrates how changes in individual glycan units can alter molecular dynamics and processing of the Env-glycan shield and, consequently, Env function.

## Introduction

The HIV-1 envelope glycoprotein (Env) is a trimer of gp160 proteins cleaved into two functional subunits: the gp41 trans-membrane glycoprotein and the exposed gp120 glycoprotein that recognizes CD4 receptor and CCR5 and/or CXCR4 co-receptors during virus-cell entry (Liu *et al*, 2008, Rizzuto *et al*, 1998, Wyatt *et al*, 1998). The multiple N-glycans of gp120 contribute to over 50% of the gp120 molecular mass (Kwong *et al*, 1998, Lee *et al*, 1992, Li *et al*, 1993, Pollakis *et al*, 2001), cover most of the Env protein surface, and are involved in key functions of HIV-1 biology (Bonsignori *et al*, 2012, Hessell *et al*, 2009, Krumm *et al*, 2016, Shivatare *et al*, 2018).

Analysis of single-genome-amplified sequences of viral RNA has identified nucleotide sequences of *env* genes from viruses responsible for productive HIV-1 infection (Abrahams *et al*, 2009, Keele *et al*, 2008), termed transmitted/founder viruses, as well as from viruses isolated during the chronic-stage of infection of the same subjects (chronic-stage viruses) (Chun *et al*, 2013, Swanstrom & Coffin, 2012). HIV-1-infected subjects produce virus-neutralizing antibodies (nAbs), beginning approximately two months after appearance of virus-specific Abs (Davis *et al*, 2009, Tomaras *et al*, 2008). Generally, transmitted/founder viruses are sensitive to autologous nAbs (Liao *et al*, 2013, McCurley *et al*, 2017), which can impose selective pressure driving the emergence of mutated immune-escape variants. Sequence analysis of such variants suggests that the escape mechanism involves mutations in potential N-glycosylation sites (NGS) of Env gp120, resulting in loss of some potential NGS and appearance of new potential NGS (Richman *et al*, 2003, Wei *et al*, 2003). Consequently, the concept of an “evolving glycan shield” has been proposed (Richman *et al*, 2003, Wei *et al*, 2003) to explain the loss of neutralizing activity of the initial antibodies against newly emerging HIV-1 variants with altered composition and/or conformation of the Env glycans. Furthermore, the glycan shield has been recognized for its ability to protect from Env-specific nAbs even when N-glycans are not evolving (Kwong *et al*, 1998, Li *et al*, 1993, Pollakis *et al*, 2001, Stewart-Jones *et al*, 2016). However, the impact of these potential NGS mutations was only studied regarding their immediate location, and escape from nAbs, but not with respect to global effects on the Env structure. Our recent analysis of all deposited HIV-1 sequences in the LANL database demonstrated that certain N-glycan microdomains have a limited number of potential NGS combinations (Hargett *et al*, 2019). These findings suggest that some glycan units have a distinct functional role in terms of position and glycan density to maintain the overall massive glycan shield (Hargett *et al*, 2019).

In this study, we took advantage of two naturally occurring HIV-1 Env variants from the same donor that differed in functional characteristics and potential NGS composition and utilized them as a model system to define how glycans can affect the structure and function of Env. By using our high-resolution glycomics analysis, we have shown that changes to key potential NGS had a ripple-effect on Env-glycan processing at distant locations, with several glycans exhibiting limited processing. These site-specific alterations of Env glycans had impact on antibody recognition and interaction with CD4 receptor. Molecular-dynamic simulations enabled visualization of the complex changes induced by specific oligosaccharide positions and indicated that some glycans are able to move to compensate for a loss of a glycan. Together, these complementary applications of molecular biology, modeling, and functional glycomics revealed how individual glycans of the massive Env glycan shield can alter the Env glycan processing and, consequently, Env function.

## Results

### Selection and characterization of HIV-1 Env with distinct glycosylation-site patterns

An essential prerequisite for studying glycan composition effects should be the identification of naturally occurring viral variants from a single donor with differing potential Env glycosylation-site composition and different functional characteristics. By using multiple-sequence alignment and NetNglyc software, we assessed WEAU HIV-1 Env from early and late stages of infection for encoded potential NGS (Fischer *et al*, 2010, Wei *et al*, 2003). We selected two Env variants with the most extreme differences in potential NGS, one from an early virus from day 16 after onset of symptoms of the acute retroviral syndrome (designated WEAU-d16) and the other from a late virus from day 391 (designated WEAU-d391) for further analyses. Env gp120 of WEAU-d16 and WEAU-d391 exhibited seven differentially positioned potential NGS in the V1, C2, the C2-V3 junction, and V4 regions (Fig 1A and B); three unique potential NGS (N141, N295, and N398) were detected in WEAU-d16 and four unique potential NGS (N234, N262, N396, and N406) in WEAU-d391.

**Figure 1.**
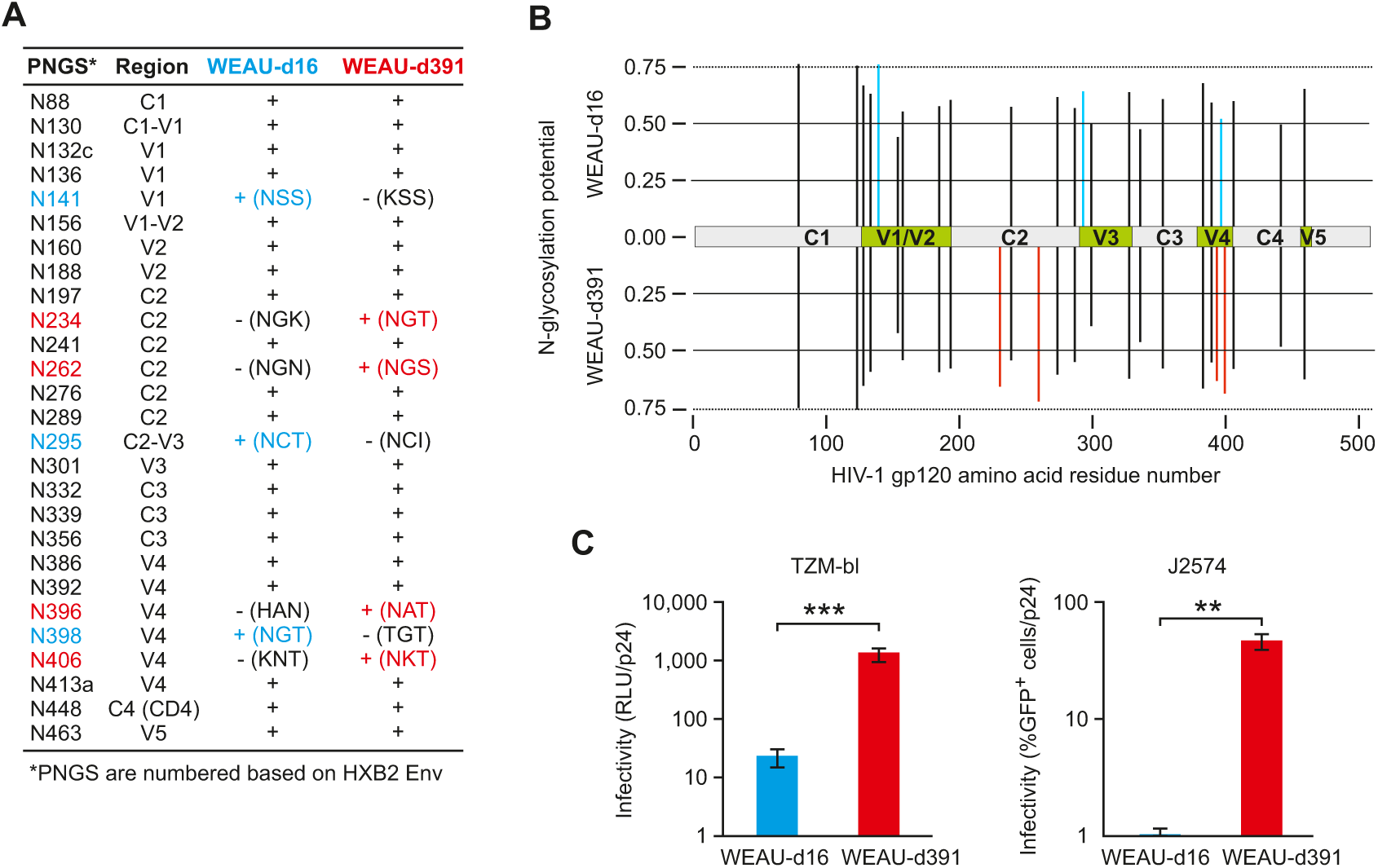
Differences in potential N-glycosylation sites (NGS) of WEAU gp120 and infectivity of WEAU early virus (WEAU-d16) and chronic-stage virus (WEAU-d391). A Potential NGS determined by amino-acid motifs N-X-S/T (N, Asn; S, Ser; T, Thr) in gp120 of WEAU-d16 and WEAU-d391. Abbreviation: PNGS, potential N-glycosylation sites; C, constant regions; V, variable regions. B *In silico* comparative analysis of WEAU-d16 and WEAU-d391 gp120 potential NGS performed by http://www.cbs.dtu.dk/services/NetNGlyc/. The threshold of likely glycosylation was set at 0.25 and the predicted glycosylation sites are marked by a vertical line. Seven unique potential NGS are marked by blue (WEAU-d16) and red (WEAU-d391) vertical lines. Constant (C) and variable (V) regions are marked. C Infectivity of WEAU Env-pseudotyped viruses produced in 293F cells using TZM-bl cells (left) or J2574 reporter T cells (right). Virus infectivity using TZM-bl reporter cells is expressed in relative light units (RLU) normalized to virus load (quantified by p24 content). Virus infectivity using J2574 reporter T cells is expressed as percentage of GFP-positive cells normalized to virus load (p24). Data information: In D, data from eight TZM-bl and three J2574 independent experiments are shown as means ± standard deviations. *** p<0.0000001; ** p<0.001.

### Functional characterization of the selected HIV-1 WEAU variants

To assess the functional impact of differential potential NGS in the selected WEAU Envs, we generated synthetic *gp160* genes of WEAU-d16 and WEAU-d391 and produced WEAU Env-pseudotyped viruses. These *gp160* genes had identical *gp41* segments (derived from WEAU-d16) whereas *gp120* genes, based on WEAU-d16 sequence, reflected the strain-specific potential NGS (Fig EV1). The infectivity of WEAU-d16 virus was significantly lower compared to WEAU-d391 using TZM-bl reporter cells (Montefiori, 2009) (Fig 1C, left; Appendix Fig S1) and J2574 reporter T-cells (Jones *et al*, 2007) (Fig 1C, right).

Differential infectivity was not the result of potential artifacts in the production of Env-pseudotyped viruses, as revealed by western blotting with serum IgG from an HIV-1-infected subject. The viral protein content and processing, including uncleaved Env (gp160), cleaved Env (gp120, gp41), p24, and other proteins, were similar for WEAU-d16 and WEAU-d391 virions (Fig 2A and B, Appendix Fig S2).

**Figure 2.**
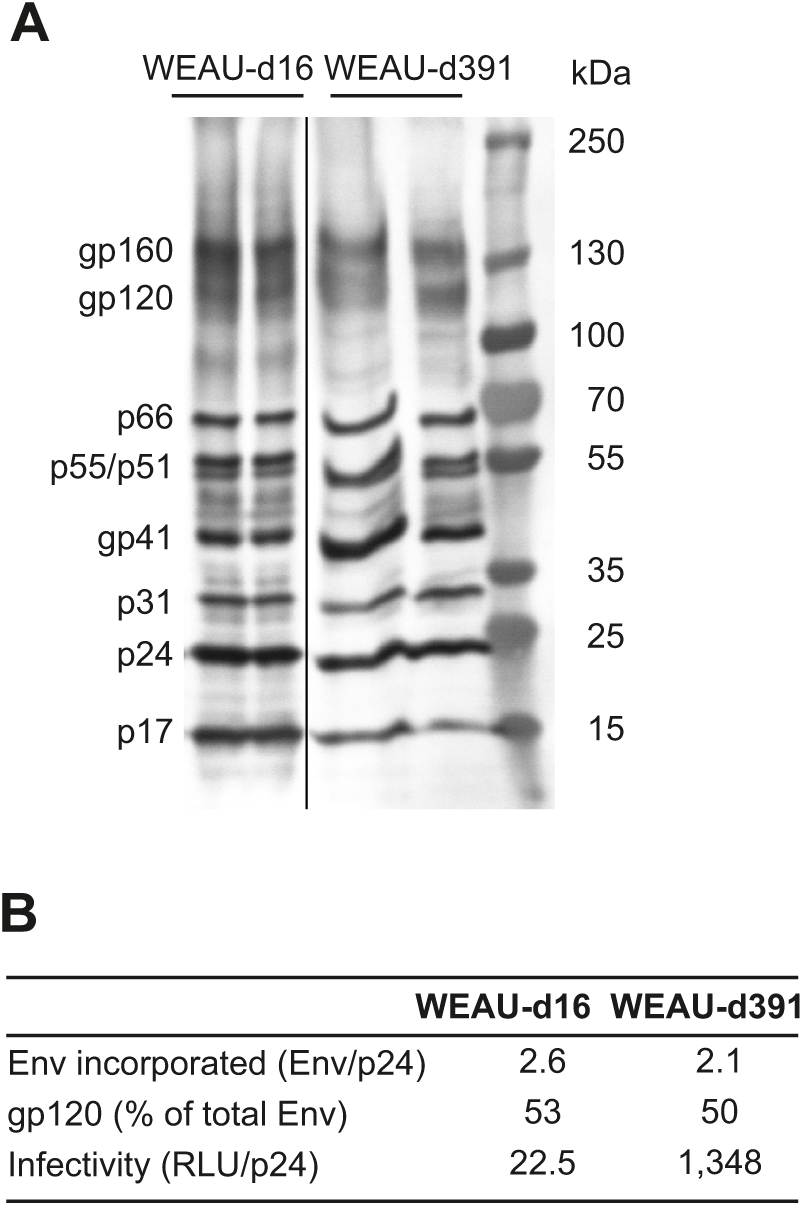
Virions with Env from an early virus (WEAU-d16) and chronic-stage virus (WEAU-d391) exhibit similar protein components, including Env expression, incorporation, and cleavage. A Representative western blot assays of 20-times concentrated viral stocks (two preparations each). Nine HIV-1 proteins were detected using serum IgG antibodies from an HIV-1-infected individual. The amounts of uncleaved Env (gp160), cleaved Env (gp120), and p24 proteins were quantified by densitometry as integrated intensity (i.i.) of each protein band. B Env incorporation into virions was calculated using the formula (gp160 i.i. + gp120 i.i.)/p24 i.i. and the Env cleavage (gp120) was calculated using the formula gp120 i.i./(gp120 i.i. + gp160 i.i.). Virus infectivity was determined in eight independent experiments using TZM-bl reporter cells and expressed in relative light units (RLU) normalized to virus load (p24).

### The N262 glycans affect binding to CD4 and recognition by CD4-binding-site-targeting broadly neutralizing antibodies in native as well as recombinant gp120 trimers

Among the seven distinct potential NGS between WEAU-d16 and WEAU-d391, we were particularly interested in the N262 NGS, the site present in WEAU-d391 but not in WEAU-d16 (Fig 1A and B). This site, previously shown to affect HIV-1 infectivity (Mathys *et al*, 2014, Wang *et al*, 2013), is not located in any Env domain with a designated direct functionality (*e*.*g*., CD4-binding site), but rather involved in gp120 folding and stability (Kong *et al*, 2015). To test whether the low infectivity of WEAU-d16 may be due to lack of the N262 glycans, we generated WEAU-d16 Env with the N262 NGS added (N264S; designated WEAU-d16-N264S) and WEAU-d391 with N262 NGS removed (S264N; designated WEAU-d391-S264N). Indeed, WEAU-d16-N264S Env-pseudotyped virus exhibited 9- and 7-fold higher infectivity in TZM-bl and J2574 reporter cells compared to the parental WEAU-d16 virus (Fig 3A and B). Conversely, WEAU-d391-S264N virus exhibited a significantly reduced infectivity compared to the parental WEAU-d391 virus; 100- or 20-fold reduced infectivity of TZM-bl or J2574 reporter cells, respectively (Fig 3A and B).

**Figure 3.**
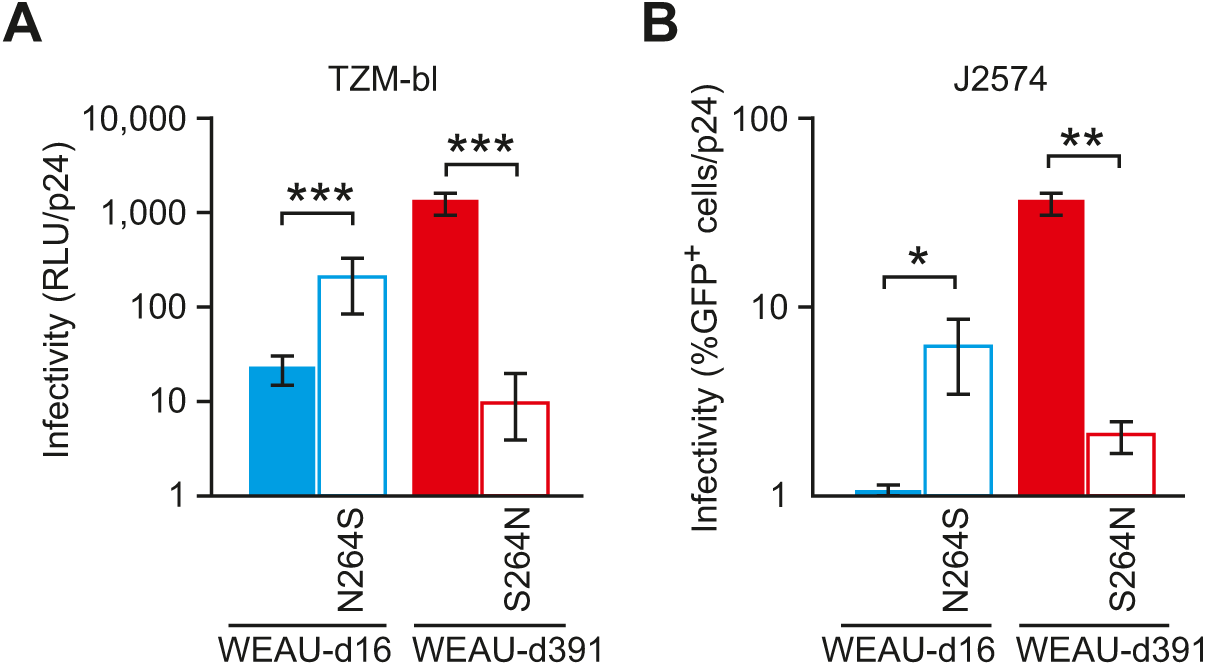
Infectivity of virions pseudotyped with Env from an early virus (WEAU-d16) and Env from chronic-stage virus (WEAU-d391) is impacted by the presence of N262 glycan. A Viral infectivity determined using TZM-bl reporter cells is expressed in relative light units (RLU) normalized to virus load (p24). B Infectivity using J2574 reporter T cells is expressed as percentage of GFP-positive cells normalized to virus load (p24). WEAU-d16-N264S is a WEAU-d16 mutant with N262 potential NGS introduced. WEAU-d391-S264N is a WEAU-d391 mutant without N262 potential NGS. The WEAU viruses without N262 potential NGS had significantly lower infectivity. Data information: in B, data from eight TZM-bl and three J2574 independent experiments are shown as means ± standard deviations. *** p<0.001; ** p<0.01; * p<0.05.

To define whether the loss of infectivity associated with the absence of the N262 NGS was related to altered virus-CD4 binding, we tested binding of WEAU Env-pseudotyped viruses to recombinant soluble CD4 (sCD4). Infectivity of viruses containing the N262 NGS was inhibited by sCD4 in a dose-dependent manner (Fig 4A and B). The IC_50_ was 2.46±0.21 µg/ml for WEAU-d391 and 1.43±0.42 µg/ml for WEAU-d16-N264S. Inhibition by sCD4 was approximately 4-fold lower (measured as IC_25_) when N262 NGS was removed from WEAU-d391 (Fig 4A and B). Similarly, inhibition by sCD4 was approximately 6-fold lower for WEAU-d16 (Env without N262 NGS) compared to the mutant with N262NGS added (WEAU-d16-N264S). These results suggest that, although N262 NGS is not located in the CD4-binding site, the N262-associated glycan controls the Env conformation in regards to CD4-binding capacity.

**Figure 4.**
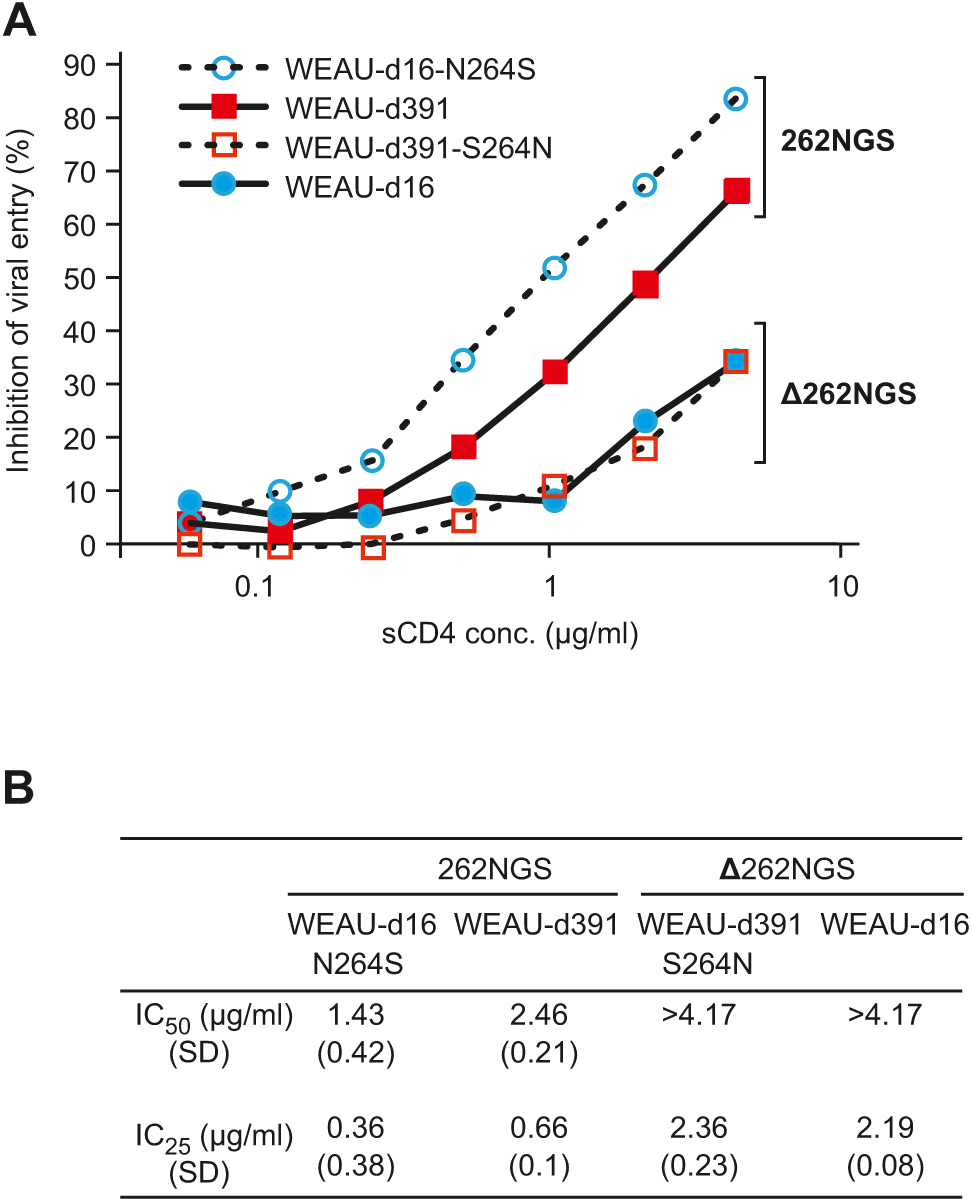
Soluble CD4 inhibits infectivity of WEAU Env-pseudotyped viruses with N262 NGS more than those without N262 NGS. A Representative results from one of three independent experiments. 262NGS indicates N262 NGS presence (WEAU-d391 and WEAU-d16-N264S), Δ262NGS indicates N262 NGS absence (WEAU-d16 and WEAU-d391-S264N). B Mean values ± standard deviations were calculated for IC_50_ and IC_25_.

To further clarify the function of the N262 glycans, we produced recombinant trimeric gp120 WEAU variants using our previously published approach that added a foldon and V5 and His tags to the gp120 (Hargett *et al*, 2019, Raska *et al*, 2014, Raska *et al*, 2008, Raska *et al*, 2010). We confirmed that gp120 WEAU glycoproteins were exclusively produced as trimers (Fig 5A).

**Figure 5.**
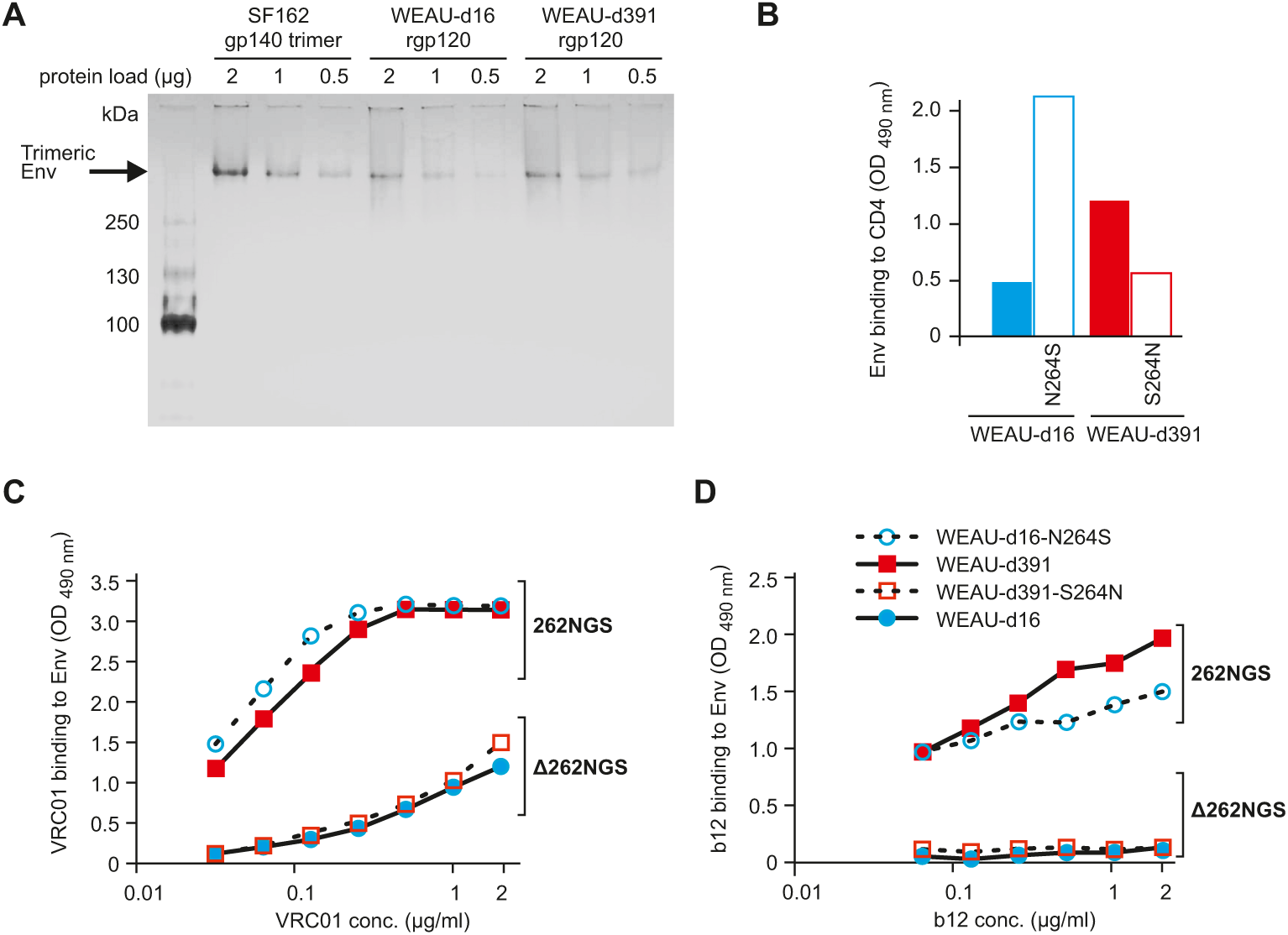
Binding of soluble CD4 and CD4-binding-site-specific broadly neutralizing antibodies (BnAbs) to recombinant gp120 trimers is enhanced by the presence of N262 glycan. A Recombinant proteins WEAU-d16, WEAU-d391, and a reference gp140 trimer (HIV-1_SF162_) (at 2, 1, and 0.5 µg protein) were fractionated using a 4-15% polyacrylamide gradient Mini-PROTEAN TGX precast gel (Bio-Rad) under native conditions. Protein bands ∼420 kDa visualized using Bio-Safe Coomassie blue G-250 stain indicated that the preparations are forming trimers under native conditions. B The binding of recombinant gp120 trimers (rgp120) to ELISA plate-coated soluble CD4 was measured as OD at 490 nm using sCD4-coated ELISA plates and expressed as OD_490_ values normalized to rgp120 protein. C,D Equal amounts of rgp120 were added to ELISA plates coated with mouse anti-His antibody. Serially diluted samples of VRC01 (B) or b12 (C) BnAb were then added and Ab binding was expressed as OD_490_ values. 262NGS indicates N262 potential NGS presence (WEAU-d391 and WEAU-d16-N264S) and Δ262NGS indicates N262 potential NGS absence (WEAU-d16 and WEAU-d391-S264N).

The four trimeric recombinant WEAU gp120 (rgp120) variants were analyzed by ELISA and western blotting to determine rgp120 protein concentrations (Raska *et al*, 2014). rgp120 WEAU-d16 and WEAU-d391-S264N exhibited low level of binding to ELISA-coated sCD4, whereas binding of rgp120 WEAU-d16-N264S and WEAU-d391 was two- to four-fold higher (Fig 5B). These results reproduced the data obtained for sCD4-mediated inhibition of virus infectivity with the respective WEAU Env variants (Fig 4).

The trimeric rgp120 variants also allowed us to test whether the N262 NGS affected recognition of the CD4-binding site by the well characterized broadly neutralizing antibodies (BnAbs) b12 and VRC01. Both BnAbs exhibited strong binding to rgp120 variants with the N262 NGS (WEAU-d16-N264S and WEAU-d391) at the range of concentrations tested (0.0625-2 µg/ml) (Fig 5C and D). Conversely, b12 did not bind to and VRC01 had a reduced ability to recognize rgp120 variants without N262 NGS (WEAU-d16 and WEAU-d391-S264N).

Together, these data demonstrate that the N262 glycans, although not located in the CD4-binding site, are crucial for Env-CD4 binding and viral infectivity. Essential for the ensuing glycomics studies, the data demonstrate that the corresponding structural and functional characteristics of native Env, as expressed on the virus, are preserved in the recombinant Env variants.

### The N262 N-glycosylation site impacts the protein-wide glycan composition of rgp120 trimer in a site-specific context

Given the effect of the N262 glycans on Env structure and functionality, we decided to decipher how the N262 glycans could globally affect the Env glycan shield. To begin to address this question, we performed monosaccharide compositional analysis by using gas-liquid chromatography to assess the global glycan composition of WEAU rgp120 trimer variants (Hargett *et al*, 2019, Raska *et al*, 2014, Raska *et al*, 2010). Surprisingly, the monosaccharide compositional analysis revealed that the presence of N262 not only affected function, but also controlled the overall glycan composition of the Env gp120 trimers. Env variants with the N262 potential NGS (WEAU-d391 and WEAU-d16-N264S) had a lower galactose content compared to Env variants without N262 potential NGS (WEAU-d16 and WEAU-d391-S264N). Galactose to mannose ratios were 0.51 and 0.42 for WEAU-d16 and WEAU-d16-N264S and 0.54 and 0.64 for WEAU-d391 and WEAU-d391-S264N. As galactose is present in most complex glycans, but absent from high-mannose glycans, this observation suggested overall enhanced glycan processing toward complex glycans in trimeric rgp120 variants without N262 potential NGS.

To determine the specific nature of changes at the individual potential NGS, we used our validated LC-MS-based glycomics workflow (Hargett *et al*, 2019). Trimeric rgp120 preparations with N262 NGS (WEAU-d16-N264S and WEAU-d391) and without N262 potential NGS (WEAU-d16 and WEAU-d391-S264N) were normalized for total protein (Fig EV2A), digested with various combinations of proteases and glycosidases, and the subsequent (glyco)peptides were analyzed by LC-MS. The glycopeptides for each NGS were expressed as relative abundance of each individual N-glycopeptide at each site for each rgp120 variant (Fig 6A). Three different types of heterogeneity were observed that categorized individual NGS as (i) predominantly high-mannose glycans, (ii) predominantly complex glycans, and (iii) a mixed population of both high-mannose and complex glycans (Fig EV2B) (Hargett *et al*, 2019). N-glycan site-specific heterogeneity profiles of the four WEAU rgp120 trimers are shown in Fig 6A (Behrens *et al*, 2017). As expected, we found N262 glycosylated in WEAU-d391 and WEAU-d16-N264S rgp120 variants, with the site occupied by a mixture of high-mannose and complex glycans.

**Figure 6.**
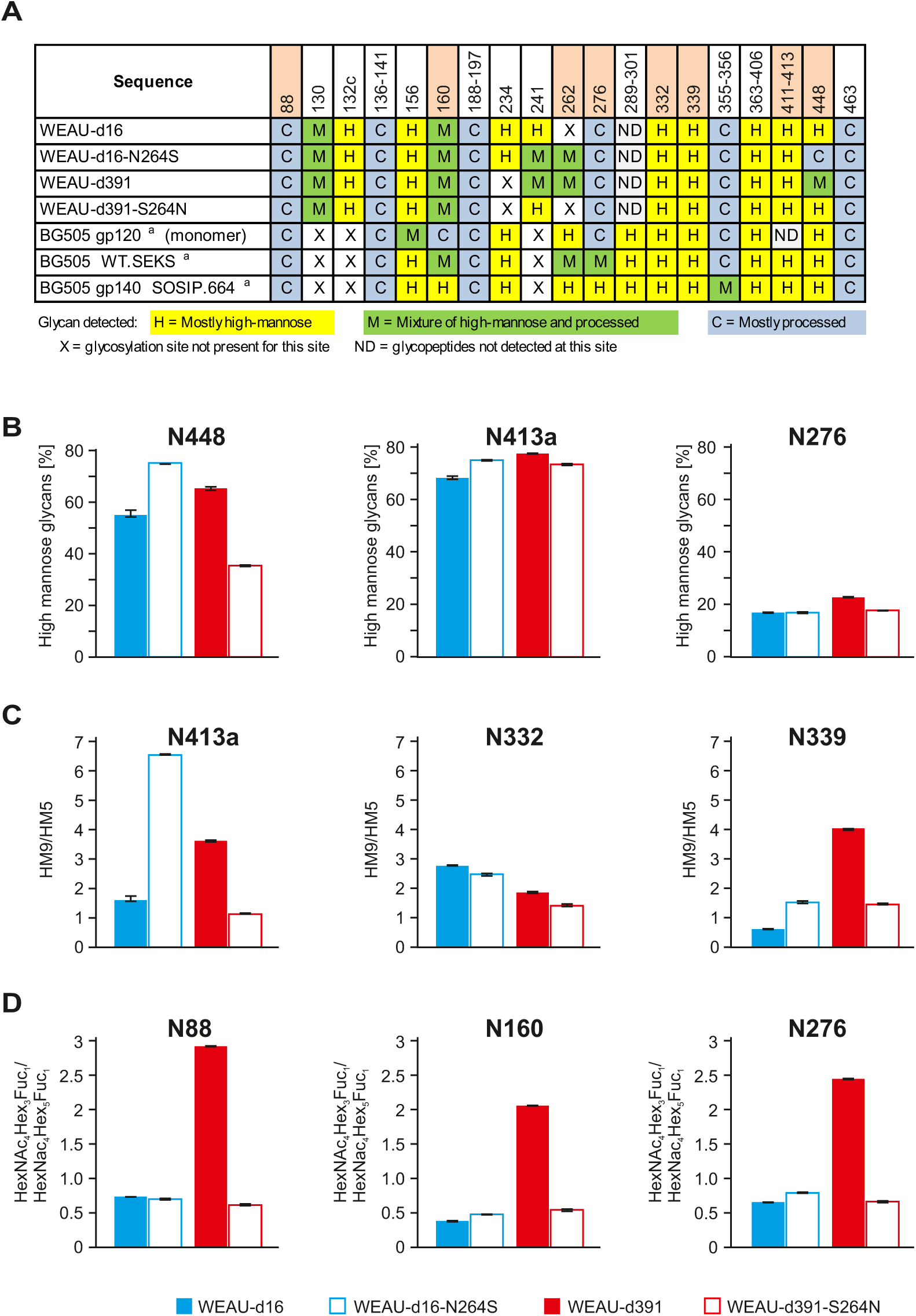
Cross-sample comparison of site-specific glycosylation heterogeneity of four different WEAU gp120 trimers reveals the impact of N262 glycan on heterogeneity of several other, spatially adjacent glycans. A Site-by-site comparison of the glycosylation profiles of four WEAU gp120 trimers (WEAU-d16, WEAU-d391, WEAU-d16-N264S, WEAU-d391-S264N) compared with those of BG505 gp120 monomer and two BG505 trimers (_a_from reference(Go *et al*, 2017)). N262 and the glycosylation sites shown in panels b-d are highlighted with pink-color background. B Comparison of abundance of high-mannose glycans at three representative NGS: N448 (variable N-glycan composition in different WEAU gp120 trimers), N413a (predominantly high-mannose N-glycans), and N276 (predominantly processed N-glycans). There was significantly more high-mannose glycans at N448 in gp120 trimers with N262 NGS. C The area under the curve of the extracted ion chromatogram (XIC) of the Man_9_ oligosaccharide (HM9) to the Man_5_ oligosaccharide (HM5) for three predominantly high-mannose-containing NGS. N413a contained significantly more Man_9_ glycans than Man_5_ in gp120 trimers with N262 NGS. N332 showed minor differences and N339 had significantly more Man_9_ glycans only in the WEAU-d391 gp120 trimer. D The ratio of area under the curve of XIC for the HexNAc_4_Hex_3_Fuc_1_ oligosaccharide to HexNAc_4_Hex_5_Fuc_1_ oligosaccharide for predominantly processed N-glycans. N88, N160, and N276 NGS had significantly more HexNAc_4_Hex_3_Fuc_1_ than HexNAc_4_Hex_5_Fuc_1_ for the WEAU-d391 gp120 trimer, implying less processing at these sites for this variant. Color coding for panels is shown at the bottom, below panel D.

To determine the effect of N262 NGS on other Env glycans, quantitative differences in the site-specific heterogeneity of individual NGS were assessed at the overall classification level (relative proportion of high-mannose, hybrid, and complex glycans) as well as at the representation of individual oligosaccharides. Of the sites that were common among all four WEAU Env variants, N448 exhibited the greatest differences in its overall N-glycan profile (Fig 6A and B). In Env variants with N262 NGS (WEAU-d16-N264S and WEAU-d391), N448 NGS had >65% high-mannose glycans. In Env variants without N262 NGS, representation of high-mannose glycans at N448 was reduced by 20% (WEAU-d16-N264S vs. WEAU-d16) and by 30% (WEAU-d391 vs. WEAU-d391-S264N). The remaining NGS exhibited relatively small differences (<10%) in high-mannose-glycan content across the WEAU variants (Fig 6A). These NGS followed the quantitative trends shown in Fig 6B for the predominantly high-mannose at N413a NGS and the predominantly processed glycans N276 NGS.

Consistent with the idea that extensive global NGS site changes will not occur as they would abolish viral infectivity, most NGS of the four WEAU rgp120 variants did not differ in their category of single-site heterogeneity (i.e., predominantly high-mannose or complex glycans, or mixture of both types), whereas several key NGS showed differences within the same category of glycans (e.g., different proportion of Man_9_GlcNAc_2_ to Man_5_GlcNAc_2_ high-mannose glycans). In Fig 6C, we demonstrated these effects by comparing the ratio of the area under the curve from the extracted ion chromatogram (XIC) of the Man_9_GlcNAc_2_ oligosaccharide to that of the Man_5_GlcNAc_2_ oligosaccharide at NGS N413a, N332, and N339. N413a NGS had more Man_9_GlcNAc_2_ than Man_5_GlcNAc_2_ in rgp120 variants with N262 NGS, indicating less processing at N413a NGS in WEAU rgp120 trimers with N262 NGS vs. those without N262 NGS. In contrast, similar ratio of these two high-mannose glycans (Man_9_GlcNAc_2_ and Man_5_GlcNAc_2_) at N332 in all four WEAU variants showed little impact of N262 NGS. Interestingly, NGS N339 in the WEAU-d391 rgp120 trimer had significantly more Man_9_GlcNAc_2_ than Man_5_GlcNAc_2_ when compared to the other three WEAU rgp120 variants (Fig 6C). A similar pattern of specific glycan abundance in the WEAU-d391 rgp120 trimer vs. other trimers was observed for the quantitative ratio of two complex glycans (HexNAc_4_Hex_3_Fuc_1_ and HexNAc_4_Hex_5_Fuc_1_) at NGS with complex glycans (N88, N160, N276; Fig 6D). This observation suggested that N-glycan shield of the WEAU-d391 variant was the least processed of the WEAU trimers analyzed, even at NGS that primarily contain complex N-glycans.

Thus, high-resolution quantitative glycomics can detail how a local, single glycosylation site (N262 NGS) influenced the N-glycan heterogeneity of rgp120 trimers and exert a glycomic ripple effect that potentially propagates functional and conformational changes throughout the Env trimer.

### Modeling the impact of positional glycosylation differences on global Env structure

The glycomic profiling detailed above provided high-positional resolution of changes imparted by N262 NGS. To begin to understand the implications of these changes, we mapped the glycoprofiling data of WEAU variants onto the structure of the gp120 trimer. The observed differences included a sub-cluster of interdependent NGS spatially close to N262 (Hargett *et al*, 2019), a so called “high-mannose patch” (Pritchard *et al*, 2015) that consists of N295, N301, N332, N413a, and N448 NGS.

We performed a series of molecular dynamics simulations (MDS) of the two WEAU variants and the corresponding N262 mutants. By using the solved crystal structure of HIV-1 clade B JR-FL prefusion Env trimer as a template (PDB: 5FYK) (Stewart-Jones *et al*, 2016), homology models were generated for all four WEAU variants (WEAU-d16, WEAU-d16-N264S, WEAU-d391, and WEAU-d391-S264N). Man_5_GlcNAc_2_ oligosaccharides were modeled at each sequon using an in-house glycosylator software. To analyze glycan motions in different WEAU trimers, 500-nanosecond (ns) MDS were performed (**Appendix Supplementary movies**). Fig 7, showing initial glycan positioning for MDS within the high-mannose patch of Env gp120, illustrates different glycan densities of the clustered glycans of this region in the four WEAU Env variants. A qualitative assessment of WEAU-d391-S264N MDS, the variant with the lowest glycan density within the high-mannose patch, revealed N301, N332, N413a, and N448 oligosaccharides to exhibit relatively free movements. Notably, the protein cleft normally occupied by glycan N262 was vacant. As the number of glycans within the high-mannose patch increased by one glycan for WEAU-d16 and WEAU-d391 or by two glycans for WEAU-d16-N264S, movement of oligosaccharides became more restricted in these trimers. These findings indicate that glycan density in the high-mannose patch differentially impacts glycan movements and that specific glycans, such as N262, can have large effects with functional implications.

**Figure 7.**
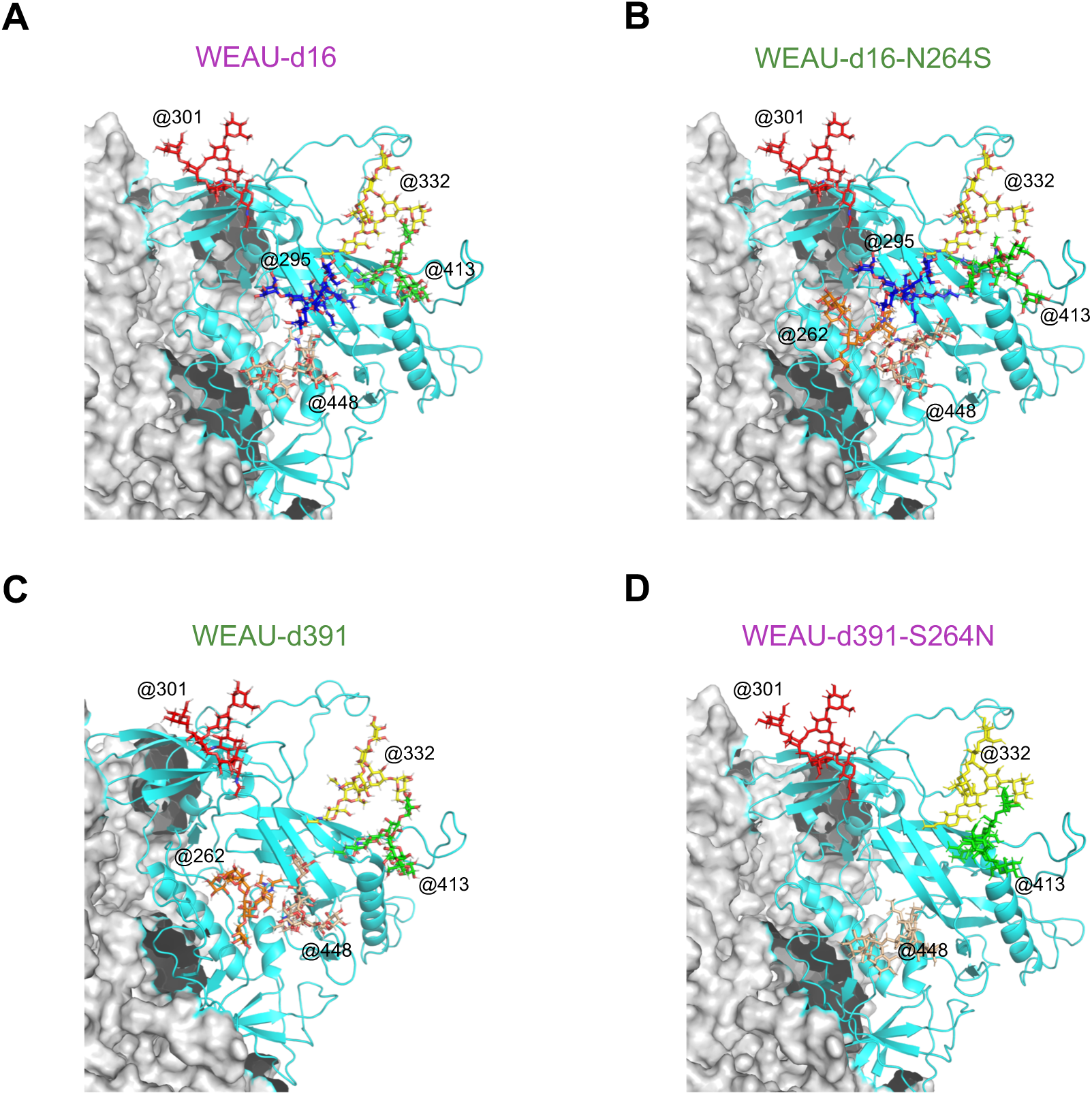
Initial glycan positioning for MDS within the high-mannose patch of envelope gp120 trimers WEAU-d16, WEAU-d16-N264S, WEAU-d391, and WEAU-d391-S264N. Glycan conformations are modeled onto the HIV-1 envelope trimer molecule using an in-house glycosylator software that arranges glycans avoiding atom clashes. Selected glycans are highlighted in stick conformation for: A WEAU-d16, B WEAU-d16-N264S, C WEAU-d391, D WEAU-d391-S264N. Glycans are marked by numbers and color-coded as in the Supplementary movies: Orange: N262 (if present), Blue: N295 (if present), Red: N301, Yellow: N332, Green: N413, Pink: N448.

Based on the initial qualitative assessment of the MDS for the four WEAU variants, we next performed a network analysis on the dense array of glycan-glycan interactions. We used changes in betweenness centrality (Kwong *et al*, 1998, Li *et al*, 1993, Pollakis *et al*, 2001, Stewart-Jones *et al*, 2016), a measure representing the number of shortest paths that pass through a node, as a means to identify changes in the interactions between the glycans. A heatmap of the betweenness centrality for each oligosaccharide was generated and overlaid onto each WEAU model (Fig EV3). When all WEAU variants were compared, we observed no differences in the betweenness centrality at the CD4-binding site. However, the extent of betweenness centrality varied among protomers at the high-mannose patch and the trimer apex. At protomer 2, there is a clear path of betweenness centrality from the base of the trimer to the apex in all WEAU variants. At protomer 3, the WEAU-d391-S264N high-mannose patch contained little or no betweenness centrality while all other variants had a high degree of betweenness centrality. At protomer 1, the high-mannose patch of WEAU-d16 and WEAU-d391-S264N contained disconnected regions of betweenness centrality while WEAU-d391 and WEAU-d16-N264S exhibited a continuous trail of glycans with a higher betweenness centrality from the apex of the trimer to the base. Thus, these features differentiated the variants with N262 glycan (WEAU-d391, WEAU-d16-N264S) from those without the N262 glycan (WEAU-d16, WEAU-d391-S264N).

Next, we averaged the location of the center of mass (COM) for each oligosaccharide spatially close to the N262 NGS over the 500-ns MDS (Figs 8 and EV4). To determine the effect the N262 glycan had on the position of the neighboring glycans, we determined the differences in the COM locations of those glycans in WEAU-d16-S264N (with N262) and WEAU-d16 (without N262) (Fig 8A). N413a shifted 12.52 Å toward N295 in WEAU-d16-S264N compared to WEAU-d16. N295 shifted 11.75 Å toward N448 in WEAU-d16-S264N. N448 shifted 7.71 Å toward N262 in WEAU-d16-S264N. N301 and N332 position changed only slightly (2.92 Å and 3.57 Å, respectively). We then determined the differences in the COM locations in WEAU-d391 (with N262) and WEAU-d391-S264N (without N262) (Fig 8B). The N448 glycan shifted 5.63 Å toward N262. N332 shifted 6.23 Å away from the β-sheet. N301 and N413a shifted only 4.24 Å and 0.81 Å respectively. Interestingly, the COM of N413, N295, and N448 glycans in WEAU-d16 shifted to compensate for the loss of the N262 oligosaccharide. Conversely, in WEAU-d391, only the COM of N448 shifted toward the COM of N262 (Fig EV4). This observation indicates that the lack of glycan density in the WEAU-d391-S264N created less steric push to fill the N262 void.

**Figure 8.**
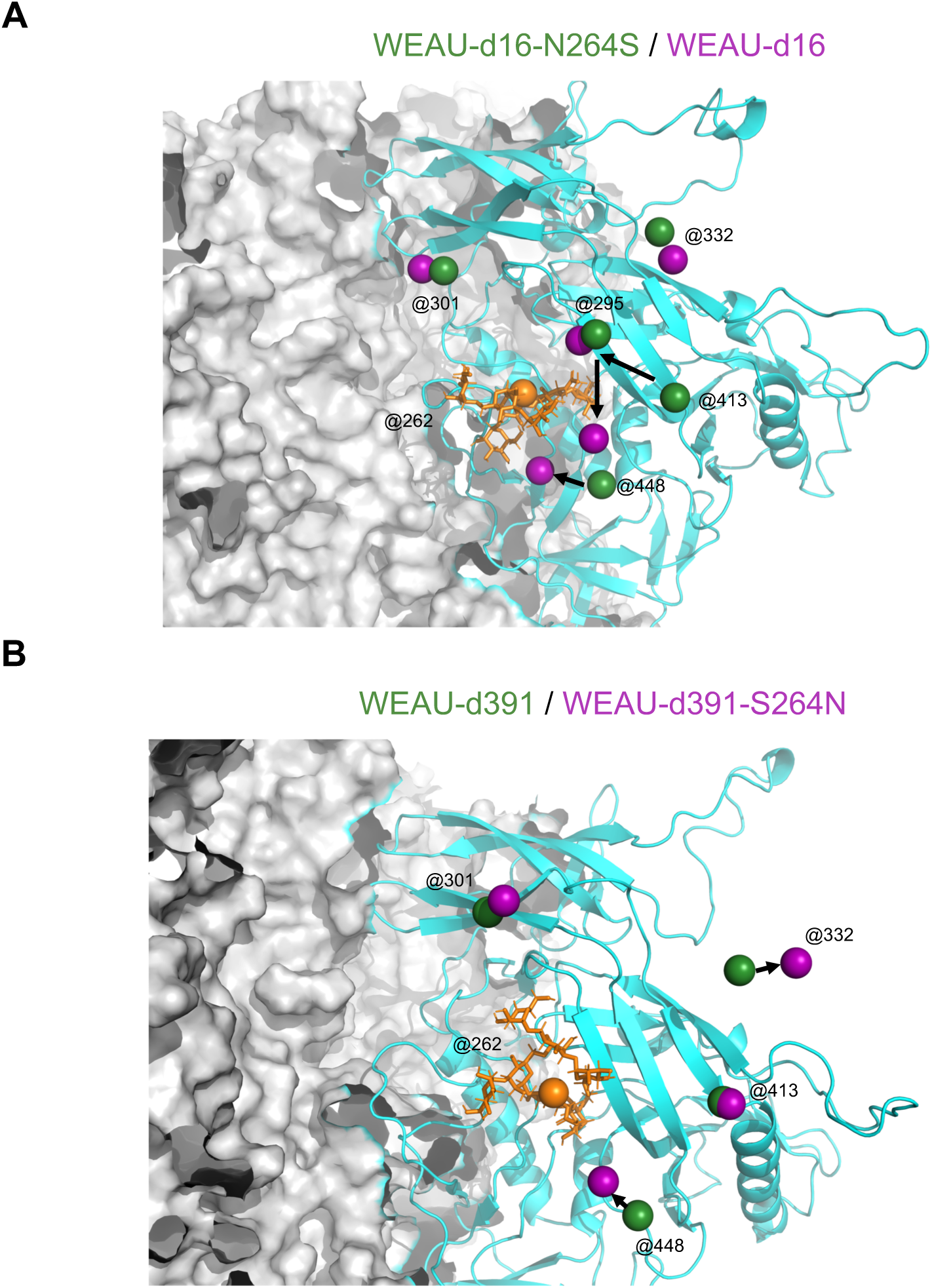
Center of Mass (COM) data for glycans within the high-mannose patch highlight glycan movement when glycan N262 is absent. A COM differences were calculated for each oligosaccharide between WEAU-d16-N264S (dark green) and WEAU-d16 (purple) at the second protomer. The resultant COM shifts were observed: 2.9 Å for N301, 3.6 Å for N332 Å, 11.8 Å for N295, 12.5 Å for N413 and 7.7 Å for N448. B COM differences were calculated for each oligosaccharide between WEAU-d391 (purple) and WEAU-d391-S264N (dark green) at the second protomer. The resultant COM shifts were observed: 4.3 Å for N301, 6.2 Å for N332 Å, 0.8 Å for N413 and 5.6 Å for N448. N262 NGS in WEAU-d16-N264S and WEAU-d391 is shown in orange.

Overall, the MDS enabled visualization of the impact of the N262 glycan on HIV-1 Env N-glycan shield. The resultant weaker glycan-glycan interactions were manifested by the altered betweenness centrality plots for WEAU-d391-S264N protomer 1 and 3 and WEAU-d16 protomer 3. Furthermore, the COM calculations revealed how specific oligosaccharide positions can move to compensate for the loss of the N262 glycan, thereby altering the N-glycan shield in this region of the Env trimer.

### N262 NGS impacts neutralization of HIV-1 by glycan-dependent/co-dependent BnAbs

Our structural data suggested several microdomains of the glycan shield of the Env trimer to be impacted by the presence or absence of N262. These microdomains are critical for the interaction of some BnAbs with Env, which provided us with an experimental means to determine the accuracy of our structural predictions. If correct, the functional implications of the observed dynamic changes in gp120 glycan shield should result in an altered ability of BnAbs to bind to the N262 Env variants. We thus tested neutralization sensitivity of HIV-1 virions pseudotyped with Env variants with N262 NGS (WEAU-d391 and WEAU-d16-N264S) and without N262 NGS (WEAU-d16 and WEAU-d391-S264N) to BnAbs with confirmed glycan-(co)dependent activity (Table 1 and Appendix Fig S3). This was indeed the case.

**Table 1.**
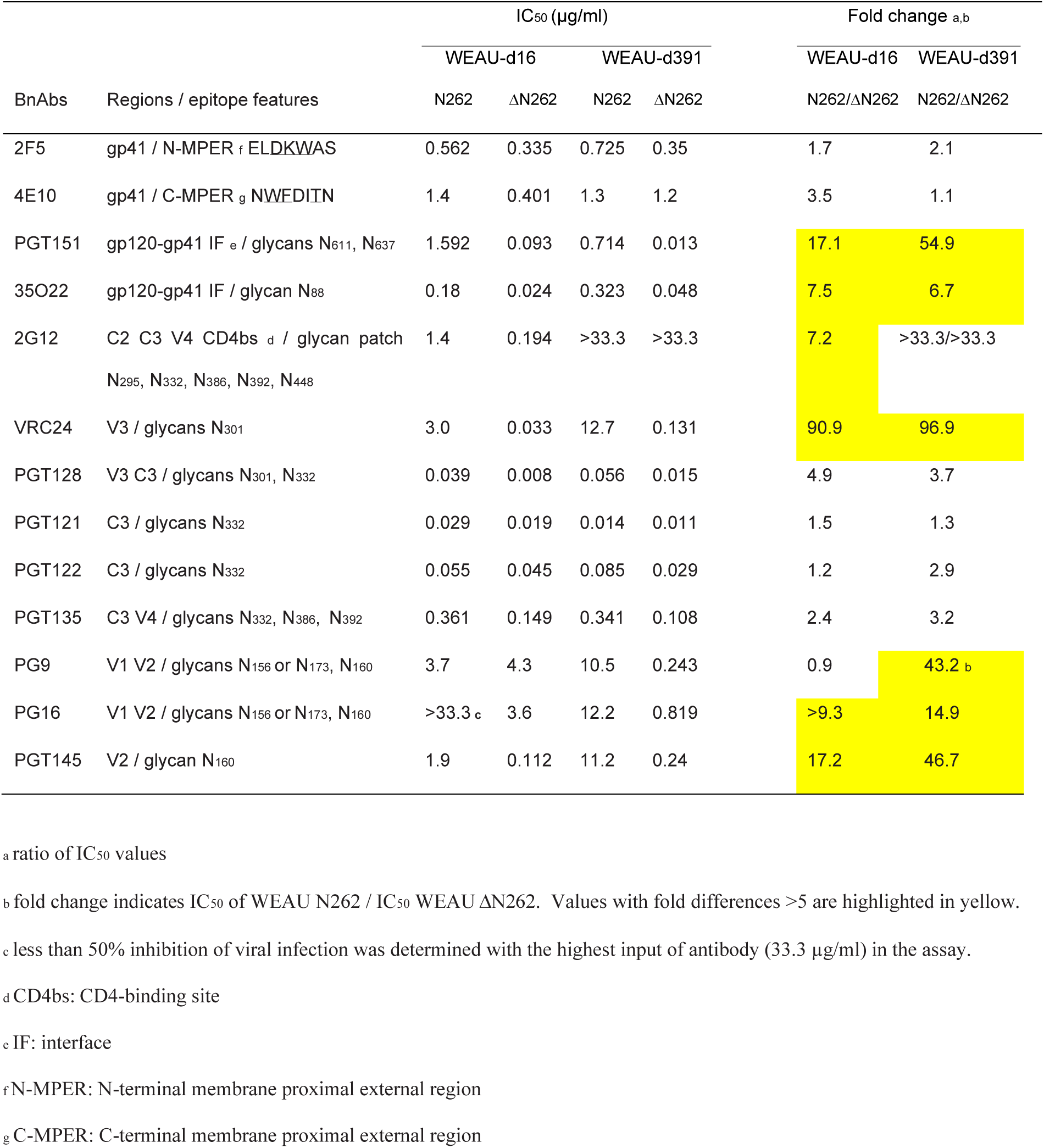
Neutralizing activity of some BnAbs against WEAU pseudotyped viruses with Env variants with and without PNGS N262 is impacted by N262 glycan

BnAbs specific for gp41 (2F5, 4E10) served as controls and showed no or only small differences in their capacity to neutralize viruses with WEAU Env variants with and without N262 NGS. In contrast, PGT151 and 35O22, BnAbs that interact within the gp120-gp41 interface (Blattner *et al*, 2014, Falkowska *et al*, 2014, Huang *et al*, 2014, Lee *et al*, 2015), exhibited differences in virus-neutralization capacity, based on N262 presence or absence. Specifically, PGT151, which has an overlapping epitope with N448, N611, N616, and N637 NGS (Lee *et al*, 2016), exhibited >50-fold better neutralization (based on IC_50_ values) of WEAU-d391-N264S compared to WEAU-d391 viruses (Table 1).

Some BnAbs with glycan-dependent binding within the V3, C3, and V4 regions of gp120 exhibited differences in neutralization activity of the tested WEAU viruses. For example, 2G12, BnAb that targets conformational epitopes consisting of high-mannose glycans (Hessell *et al*, 2009, Murin *et al*, 2014), neutralized only WEAU-d16 and WEAU-d16-N264S but not WEAU-d391 and WEAU-d391-S264N. This resistance of the immune-escape WEAU-d391 virus to 2G12 neutralization was likely due to the absence of N295 potential NGS, one of the glycans forming 2G12 epitope (Wei *et al*, 2003). WEAU-d16-N264S was 7-fold less sensitive to neutralization by 2G12 than WEAU-d16, based on IC_50_ values. Notably, serum IgG from WEAU subject five years after acquiring the infection still contained nAbs against early virus variant (WEAU-d16) but not the chronic-stage immune-escape variant (WEAU-d391) that was also resistant to BnAb 2G12 (Appendix Fig S3). VRC24, BnAb interacting with V3 and N301 NGS (Georgiev *et al*, 2013), exhibited >90-fold better neutralization (based on IC_50_ values) of WEAU virions without N262 NGS than those with N262 NGS.

Conversely, PGT128, BnAb targeting the epitope in the V3 loop and the N-glycans at N301 and N332 (Doores *et al*, 2015, Pejchal *et al*, 2011), exhibited similar neutralization capacity against all four WEAU variants. This observation is in agreement with the previous finding that for Env variants with N332 NGS, the N301 glycan becomes nonessential for neutralization by PGT128-type BnAbs (Krumm *et al*, 2016). As expected, other BnAbs that interact with N332 glycan (Krumm *et al*, 2016), such as PGT121, PGT122, and PGT135, also exhibited similar neutralization capacity against the tested WEAU viruses irrespective of N262 NGS.

PG9, PG16, and PGT145, BnAbs that target the apex of the Env trimers, exhibited differences in neutralization of WEAU pseudotyped viruses dependent on N262 NGS. PG9 and PG16 recognize glycan*-*containing regions of V1/V2, including glycans at N156 or N173 and N160 (Euler *et al*, 2011, McLellan *et al*, 2011). PG9 exhibited >40-fold higher neutralization capacity for WEAU-d391 without vs. with N262 NGS. In contrast, similar neutralization capacity was observed for WEAU-d16 variants irrespective of N262 NGS. However, both PG16 and PGT145 exhibited higher neutralization capacity for WEAU variants without N262 NGS.

In summary, as predicted by the structural modeling of the glycomics data, the neutralizing capacity of some BnAbs with glycan-(co)dependent epitopes ranging from the base of the trimer to its apex was impacted by N262 glycan. Specifically, PGT151 and 35O22 (gp120/gp41 interface), 2G12 and VRC24 (V3, C3, and V4 regions), and PG16 and PGT145 (trimer apex) exhibit reduced neutralization capacity against Env-pseudotyped viruses with N262 NGS. Together, these results confirm our glycomics and structural modeling data that suggest that the local N262 glycan has a ripple effect on the Env glycan shield structure and the Env conformation with functional implications for virion infectivity.

## Discussion

Results in this study provide new insights into how changes in glycosylation sites of the HIV-1 Env, such as loss of N262, can have ripple effect on the glycan shield with profound implications for Env structure and function. We have used two naturally occurring Env variants with different potential NGS as a model system to study how glycans affect Env structure and function. High-resolution quantitative glycomics analysis revealed that N262 NGS affected composition of several glycans at distant locations, by restricting their processing, showing how N262 and other glycans in the proximity work together. Molecular modeling and MDS revealed that some of the affected glycans are spatially close to N262, in the so called high-mannose patch, and showed that some of these glycans can move to compensate for a loss of a glycan. Other affected glycans were in the apex of the trimer. These site-specific alterations of Env glycans have significant consequences, as they impact recognition by BnAbs and interaction with CD4 receptor.

By using Env with a naturally occurring mutation at the N262 NGS (WEAU-d16), we have resolved a previously encountered problem, wherein an artificial removal of N262 NGS led to Env lysosomal degradation and, thus, production of noninfectious virions (Mathys *et al*, 2014, Wang *et al*, 2013). HIV-1 Env with naturally absent N262 NGS are rare; analysis of a database of 6,223 HIV-1 Env sequences revealed that N262 potential NGS was in 99.65% of all sequences (Hargett *et al*, 2019). In this study, we used Env from an early virus (WEAU-d16) naturally without N262 NGS and introduced the N262 NGS abrogating mutation into the matching chronic-stage virus (WEAU-d391) to generate a variant without N262 NGS (WEAU-d391-S264N). For comparison, we also engineered a WEAU-d16 variant (WEAU-d16-N264S) with added N262 NGS.

The first step in HIV-1 host-cell entry is the interaction between Env and CD4 receptor (Maddon *et al*, 1986, McDougal *et al*, 1986). Our experiments showed that the N262 glycan influenced Env-CD4 receptor interaction, further extending previous studies (Mathys *et al*, 2014, Wang *et al*, 2013). Specifically, Env-pseudotyped viruses with the N262 glycan (WEAU-d391, WEAU-d16-N264S) had higher infectivity and were more sensitive to inhibition of viral entry by sCD4 than variants without N262 NGS (WEAU-d16, WEAU-d391-S264N). Recombinant trimeric gp120 was affected in the same manner: the variants with N262 glycan exhibited higher binding to CD4 compared to the variants without N262 NGS. As N262 NGS is not a part of CD4-binding site, we speculated that the observed differences in CD4 binding represent a ripple effect of N262 glycan. Possible mechanisms may include (i) destabilization of the Env trimer due to the loss of glycan and side-chain interactions with amino-acid residues within β22 strand, demonstrated by crystallographic and electron-microscopy data (Kong *et al*, 2015, Ozorowski *et al*, 2017); (ii) an increase in Env trimer flexibility through fewer glycan-glycan interactions within the N-glycan shield, manifested by a lower betweenness centrality within protomers and an increase in processed N-glycans; and (iii) partial occlusion by N-glycans spatially close to the CD4-binding site, such as N276, which has relatively more HexNAc_4_Hex_5_Fuc_1_ glycan than HexNAc_4_Hex_3_Fuc_1_ glycan, as shown for WEAU-d16 vs. WEAU-d391 Env.

A published structure of fully glycosylated HIV-1 gp120 core revealed the N262 glycan to be buried within a weakly negatively charged protein cleft bridging the inner and outer domains of gp120 (Kong *et al*, 2015). Upon CD4 binding, the D1 arm of the N262 glycan moves by approximately 5Å into a new pocket, stabilizing an alpha helix in the CD4-bound state (Kong *et al*, 2015, Ozorowski *et al*, 2017). A loss of the N262 glycan may destabilize the gp120 core, thus affecting the affinity of CD4-binding pocket. In fact, our COM calculations from the MDS of WEAU Env trimer variants have suggested that, in the absence of N262, other oligosaccharides, such as N448, shift toward this cleft, partially complementing the role of N262 glycan.

Our glycomics analyses revealed an overall increase in processed oligosaccharides in gp120 trimers without N262, especially at NGS in the vicinity of N262 (Hargett *et al*, 2019). Even though the CD4-binding pocket is highly conserved and only peripherally surrounded by oligosaccharides (Kwong *et al*, 1998), structural studies reveal an impact of glycans, such as N276, spatially close to the CD4-binding pocket (Ozorowski *et al*, 2017, Stewart-Jones *et al*, 2016). In fact, MDS of the BG505 SOSIP Env trimer suggested that only one CD4-binding pocket is accessible at a time, due to oligosaccharides occluding the pocket on other protomers (Lemmin *et al*, 2017). Notably, WEAU-d391, the Env that yields the most infective virions among all tested WEAU variants, had the least processed N276 N-glycans. Thus, we can hypothesize that the extent of glycan processing at NGS spatially close to the CD4-binding pocket can influence accessibility of CD4-binding site. This conclusion is consistent with results of a recent MDS study that showed that in the absence of N301 glycan, another glycan that shields V3 loop and CD4-binding-site epitopes, several other glycans can structurally rearrange to maintain the glycan shield (Ferreira *et al*, 2018). Based on our recent analysis of combinations of glycans in different regions of Env trimer, we propose that some glycan microdomains would only have a limited number of NGS combinations that maintain the glycan shield (Hargett *et al*, 2019). The functional data and MDS analyses in this study corroborate this idea and further imply that variations within a glycan-microdomain density can alter Env function. Importantly, these aspects can now be effectively characterized by tracking differences in N-glycan heterogeneity profiles with site-specific resolution across Env variants, as demonstrated here.

Furthermore, our data have also revealed that the N262 NGS impacts neutralization sensitivity to some BnAbs. We observed better neutralization by PGT151, 35O22, VRC24, 2G12, VRC24, PG9, PG16, and PGT145 BnAbs for virions pseudotyped with Env without N262. Notably, the absence of the N262 glycan had little effect on BnAbs PGT128, PGT121, PGT122, and PGT135 that all recognize an epitope within the high-mannose patch that involves N332 NGS (Table 1). These observations are consistent with the glycomics profiling (Fig 6) and MDS COM calculations (Fig 8), which have shown that N332 has a similar glycan profile and COM location in all WEAU variants.

In summary, this study revealed how a naturally occurring N-glycan mutation can affect function of HIV-1 Env through altered glycosylation. Although there are many other Env variants impacting N-glycan shield to be explored, this study provides new insight into complex effects of Env glycans on HIV biology. By utilizing molecular experiments, structural analysis, and glycomics profiling, we can begin to understand the limitations of the HIV-1 “balancing act” between the viral immune escape and infectivity.

## Materials and Methods

### Reagents

All chemicals, unless otherwise specified, were purchased from Sigma (St. Louis, MO). Tissue-culture media and media-supplement were purchased from Invitrogen (Carlsbad, CA).

### Cells

FreeStyle 293-F cells (293F, Invitrogen) derived from human embryonic kidney cells (HEK) were cultured in serum-free FreeStyle 293 expression medium at 37°C in a humidified atmosphere with 8% CO_2_ on an orbital shaker platform rotating at 150 rpm. 293F were subcultured at 1:10 dilution when the density reached between 1.5 × 10^6^ and 3 × 10^6^ viable cells per ml. TZM-bl is a genetically engineered cell line derived from HeLa cells; TZM-bl cells express CD4, CXCR4, and CCR5 and contain Tat-responsive reporter genes for firefly luciferase (Luc) under regulatory control of an HIV-1 long terminal repeat (LTR) (Sarzotti-Kelsoe *et al*, 2014). 293T/17 cells derived from HEK 293T cell line are commonly used to produce high titers of infectious retrovirus. TZM-bl cell line (NIH AIDS Research and Reference Reagent Program, Catalog number 8129) and HEK cell line 293T/17 (ATCC CRL-11268) were maintained in Dulbecco’s modified Eagle’s medium (DMEM) supplemented with 10% heat-inactivated fetal bovine serum (FBS), L-glutamine (2 mM), penicillin (100 U/ml), and streptomycin (100 µg/ml) at 37°C in a humidified atmosphere with 5% CO_2_. When cell monolayers reached confluency, they were treated with 0.25% trypsin and 1 mM EDTA solution, and subcultured at 1:10 dilution. J2574 GFP-reporter T cells were generated by retrovirus transduction of Jurkat T cells (CD4^+^, CCR5^+^, and CXCR4^+^) with HIV-1 NL4.3-based LTR-GFP-LTR (p2574) construct (Duverger *et al*, 2013). The cells were maintained in complete RPMI 1640 medium (RPMI 1640 supplemented with 10% heat-inactivated FBS, L-glutamine (2 mM), penicillin (100 U/ml) and streptomycin (100 µg/ml)) and were subcultured when the density reached 1 × 10^6^ cells per ml. We used a population of J2574 T cells, representing greater than 100,000 diverse founder cells, to avoid potential clonal effects that could affect HIV-1 infection (Duverger *et al*, 2013).

### Recombinant proteins and antibodies

The recombinant soluble human CD4 (sCD4) and HIV-1_SF162_ gp140 trimer were obtained through the NIH AIDS Research and Reference Reagent Program (AIDS RRRP), Division of AIDS, NIAID, NIH. The following HIV-1 broadly neutralizing antibodies (BnAbs) were also obtained through the NIH AIDS RRRP: 2G12, PG9, PG16, VRC01, NIH45-46_G54W_ IgG, VRC03, b12, PGT121, 4E10, 2F5. Other HIV-1 BnAbs, PGT122, PGT128, PGT135, PGT145, PGT151, VRC024, 35O22, and CH01 were provided by Dr. Kwong (Vaccine Research Center, NIAID, NIH, Bethesda, MD). Plasma and serum samples from HIV-1-infected patients were obtained from the AIDS 1917 clinic, University of Alabama at Birmingham (UAB), according to an IRB-approved protocol.

### HIV-1 *env* gp160 and gp120 constructs and mutagenesis

WEAU-d16 gp160 DNA (GenBank accession number AY223743) corresponds to the WEAU16-02 single-genome-amplified sequence of viral RNA isolated from plasma HIV-1 RNA obtained 16 days after onset of symptoms of the acute retroviral syndrome (Keele *et al*, 2008, Wei *et al*, 2003). WEAU-d391 gp160 (GenBank accession number KU664677) is based on the amino-acid backbone of WEAU-d16, but the PNGS were mutated to mimic the chronic-stage WEAU391-03 gp160 from plasma HIV-1 RNA collected 391 days after onset of symptoms. Both DNA sequences coding for the respective gp160 protein were codon-optimized, synthesized by commercial supplier (ATG:biosynthetics, Merzhausen, Germany), and cloned in frame into expression plasmid pcDNA3.1 (yielding pcDNA3.1-WEAU gp160 clones that were used for production of Env-pseudotyped HIV-1). Region corresponding to gp120 amino-acid sequences shown in Fig EV1 are numbered according to the alignment with HIV-1_HXB2_ sequence (GenBank accession number K03455). The Env mutants at amino-acid position 264, designated as WEAU-d16-N264S and WEAU-d391-S264N, were prepared by site-directed mutagenesis using proof-reading platinum Pfx DNA polymerase and the following primers (sequences shown from 5’to 3’ end; letters in bold and italic indicate mutated nucleotides):

WEAU-d16 forward CTGCTGCTGAACGGCA***G***CCTGGCCGAG;

WEAU-d16 reverse CTCGGCCAGG***C***TGCCGTTCAGCAGCAG;

WEAU-d391 forward CTGCTGCTGAACGGCA***A***CCTGGCCGAG;

WEAU-d391 reverse CTCGGCCAGG***T***TGCCGTTCAGCAGCAG,

For expression of recombinant Env proteins, gp120 DNA of WEAU-d16, WEAU-d391, WEAU-d16-N264S, or WEAU-d391-S264N was fused at the 5’ end with cDNA encoding for the first 62 amino acids of non-glycosylated fragment of human mannan-binding lectin (MBL, GenBank accession number EU596574 fragment 66-252), a foldon to drive gp120 trimerization and secretion, as described before (Hargett *et al*, 2019, Raska *et al*, 2010). The resultant constructs of WEAU gp120 and MBL DNA were then cloned into pOptiVec/V5-His (designated pOptiVec/V5-His WEAU gp120-trimer) for expression and purification of trimeric gp120 (Raska *et al*, 2010). The entire *env* gene of each plasmid construct was sequenced to confirm the sequence. Plasmid DNA used in transfection of cells was purified using the Qiagen MaxiPrep kit (Qiagen, Valencia, CA).

### Expression and purification of recombinant gp120

293F cells were transfected by plasmids pOptiVec/V5-His WEAU gp120-trimer using 293fectin according to the manufacturer’s (Invitrogen) instructions. Briefly, plasmid DNA and 293fectins were mixed at the ratio of 1:2 (1 µg of plasmid DNA to 2 µl of 293fectin) and after 20-min incubation at room temperature, DNA-293fectin complexes were added to the cells at a final density of 1 × 10^6^ viable cells per ml. After 4 days, cell culture supernatants were collected, fresh serum-free FreeStyle 293 expression medium was added to the cells, and supernatants were collected again on day 7 post-transfection. The presence of recombinant gp120 (rgp120) was detected in the supernatants by SDS-PAGE and western blotting using horseradish-peroxidase (HRP)-conjugated anti-V5 antibody (see section of SDS-PAGE and western blotting for details). Then, the protein was isolated on nickel-nitrilotriacetic acid (Ni-NTA)-agarose under native conditions, according to the manufacturer’s (Qiagen) instructions. The elution buffer with rgp120 was exchanged for phosphate-buffered saline (PBS) using Amicon Ultra-4 Centrifugal Filter Unit (Millipore, Billerica, MA). The concentration of rgp120 protein was calculated by dividing optical density at 280 nm (NanoDrop 2000/2000c) by 1.21, the molar extinction coefficient determined for WEAU gp120. For normalization of rgp120 preparations, we used ELISA (see rgp120 ELISA) and western blotting after removal of all N-glycans (see the section on treatment with glycosidases for details). All rgp120 proteins were kept frozen at -80°C until use.

### Enzyme-linked immunosorbent assays (ELISAs)

For rgp120 normalization of the recombinant envelope preparations, ELISA was performed as follows: 96-well MaxiSorp plates (NUNC, Roskilde, Denmark) were coated with 100 ng/well of mouse anti-Penta His BSA-free antibody (Qiagen) in PBS, pH7.4, and incubated overnight at 4°C. After washing with PBS containing 0.05% Tween-20 (PBST), the wells were blocked with SuperBlock blocking buffer in Tris-buffered saline (TBS) (ThermoFisher Scientific, San Jose, CA) containing 0.05% Tween-20 (TBST) for 1 h at room temperature. Duplicates of serial dilution of rgp120 samples and a reference rpg120 were added to individual wells and incubated for 1 h at 37°C. The wells were washed with PBST and incubated with HRP-conjugated anti-V5 antibody (Invitrogen) at the dilution of 1:7,000. After washing, the plates were incubated with peroxidase substrate (o-phenylenediamine-H_2_O_2_) for up to 40 min at room temperature. The color development was stopped with 1 M sulfuric acid and the optical densities were measured at 490 nm (OD_490_) with an ELISA reader. The range of dilutions that yielded OD_490_ from 1 to 2 was identified and used in the following sCD4 binding assay.

To determine the relative binding affinity of sCD4 to WEAU rgp120, 96-well MaxiSorp plates were coated with 25 ng of sCD4 in 50 mM carbonate buffer, pH 9.6 and incubated overnight at 4_o_C. Duplicates of two-fold serial dilutions of rgp120 samples (concentrations normalized by ELISA) were added to individual wells and the captured Env proteins were detected with HRP-anti-V5 antibody (diluted 1:7,000) followed by the peroxidase substrate, o-phenylenediamine-H_2_O_2_. The color reaction was stopped with 1 M sulfuric acid and the optical densities were read at 490 nm.

To evaluate the binding of BnAbs to rgp120, the ELISA-normalized rgp120 samples (25 ng per well) were applied onto 96-well MaxiSorp plates coated with 100 ng/well of mouse anti-Penta His BSA-free antibody. The wells were blocked with SuperBlock in TBST for 1 h at room temperature. Duplicates of two-fold serial dilutions of BnAbs were added and incubated for 1 h at 37°C. After washing away the unbound BnAbs with PBST, the wells were incubated for 2 h at 37°C with HRP-mouse anti-human IgG antibody (dilution 1:2,000) (Southern Biotech, Birmingham, AL) followed by the peroxidase substrate, o-phenylenediamine-H_2_O_2_. The color reaction was stopped with 1 M sulfuric acid and the absorbance measured at 490 nm.

### Generation of Env-pseudotyped viruses

WEAU Env-pseudotyped viral stocks were generated by transfecting 10 µg of an *env*-deficient HIV-1 backbone vector (pSG3^ΔEnv^) and 5 µg of a WEAU gp160 plasmid (pcDNA3.1-WEAU gp160) into exponentially dividing 293T/17 cells using FuGENE 6 (Roche Applied Science, Indianapolis, IN) or into 293F cells using 293fectin (Invitrogen) according to manufacturer’s instructions. Virus-containing cell-culture supernatants were harvested 60 h post-transfection, centrifuged at 3,000 rpm for 10 min, passed through 0.22-µm filter, and kept frozen at -80°C until use. Cell-culture supernatants of transfected cells by a single plasmid pSG3^ΔEnv^ (10 µg) or WEAU env-encoding plasmid (5 µg) served as negative controls. The concentration of HIV-1 Gag p24 antigen in supernatants was determined with HIV-1 p24 ELISA kit (PerkinElmer, Inc. Waltham, MA). To analyze HIV-1 Env expression, incorporation, and cleavage (the amount of gp120 relative to that of unprocessed gp160), WEAU Env-pseudotyped viruses produced by 293F were concentrated 20-fold using Lenti-X concentrator (Clontech Laboratories, Inc. Mountain View, CA). Briefly, three volumes of prepared virus stock were mixed with one volume of Lenti-X concentrator and after 4-h incubation at 4°C, the mixture was centrifuged at 1,500 g for 45 min at 4°C. The pellet was resuspended in DPBS at 1/20 of the original culture-supernatant volume. The concentrated virus stock was stored at -80°C in single-use aliquots.

### Infectivity assay

The infectivity of WEAU Env-pseudotyped viruses was measured in a single-round entry assay by incubation of viruses with TZM-bl reporter/indicator cells in a 96-well format, using standard protocols described previously (Sarzotti-Kelsoe *et al*, 2014). To quantify the virus infectivity, the mean value of relative light units (RLU) from duplicate wells were calculated and expressed as normalized values per pg of HIV-1 p24 (RLU/p24). The dilution of a virus stock that yielded RLU between 10,000 and 100,000 was identified and used in neutralization assays, as detailed below.

The infectivity of WEAU Env-pseudotyped viruses was also measured using J2574 GFP-reporter T cells (Duverger *et al*, 2013). Serial two-fold dilutions of virus stock in 10% RPMI were incubated with J2574 (in triplicates for each dilution) in a 96-well plate for 48 h at 37°C. The infectivity of WEAU Env-pseudotyped viruses was determined by monitoring the GFP expression of J2574 cells by using a Guava EasyCyte flow cytometer (Guava Technologies, Inc.). Data analysis was performed using Guava Express software (Guava Technologies, Inc.). The infectivity of Env-pseudotyped viruses was expressed as %GFP^+^ cells per ng of HIV-1 p24 (%GFP^+^ cells/p24).

### Neutralization and inhibition assays

Neutralization activity of HIV-1-specific antibodies was measured as a reduction in Luc reporter gene expression after a single round of infection of TZM-bl cells with Env-pseudotyped viruses, as previously reported (Wei *et al*, 2012). Briefly, freshly trypsinized cells were cultured overnight in 96-well flat-bottom culture plates at a density of 10^4^ cells per well. Serial dilutions of serum/plasma samples from HIV-1-infected individuals (three-fold dilutions) and BnAbs (two-fold dilutions) were prepared in 120 µl of 10% DMEM in separate Nunc 96-well plates. The virus stocks were diluted (1:240 for WEAU-d391; 1:120 for WEAU-d16-N264S; 1:40 for WEAU-d16 and WEAU-d391-S264N) in 120 µl of 10% DMEM with 40 µg/ml of DEAE dextran (final concentration of DEAE dextran was 20 µg/ml) and added to each well of the Nunc 96-well plates, yielding a total volume in each well of 240 µl. Two sets of control wells were also included (each with at least six wells). One set contained only virus (virus control), and the other set contained medium without virus (background control). The virus-antibody and control mixtures were pre-incubated at 37°C for 1 h. After the culture medium was removed from the plates with cells, 100 µl of pre-incubated virus-antibody and control mixtures were transferred into the corresponding wells (duplicates for each dilution). After a 48-h incubation at 37°C and 5% CO_2_ incubator, the luciferase activity was measured as described above for virus infectivity assay. The reduction of virus infectivity was calculated as percentage of the average RLU in wells cultured with virus alone after the background RLU (cells only) was subtracted. The highest dilution of a serum/plasma or the lowest concentration of a BnAb that inhibited virus infection by 50% was considered the neutralization titer (IC_50_).

The standard neutralization protocol described above was also used to determine the capacity of sCD4 (1 µg/µl) to inhibit the infectivity of WEAU Env-pseudotyped viruses. sCD4 was serially diluted (two-fold dilutions), incubated with the virus, and sCD4-virus mixtures were then used to infect TZM-bl cells. The inhibition by sCD4 was evaluated as the lowest concentration of sCD4 to reduce 25% (IC_25_) or 50% (IC_50_) of virus infectivity.

### SDS-PAGE and Western Blotting

Twenty-fold concentrated viral stocks or purified rgp120 were separated by SDS-PAGE under reducing conditions using a 4-15% Mini-PROTEAN TGX precast gel (Bio-Rad Laboratory Inc. Hercules, CA), and then transferred onto a polyvinylidene difluoride (PVDF) membrane (Millipore) using Semi-Dry blotter (Bio-Rad). The membranes were blocked overnight at 4°C using SuperBlock in TBST. In addition, 1% goat serum was included in blocking buffer when goat antibodies were to be used. The membranes with blotted viral proteins were incubated with the indicated primary antibodies: IgG in sera of HIV-1-infected individuals (1:500), or 4E10 (0.3 µg/ml) or 2F5 (0.25 µg/ml) monoclonal antibodies overnight at 4°C. After being washed with PBST, the membranes were developed in sequence with biotinylated F(ab’)^2^ fragments of goat anti-human IgG (Geneway Biotech. Inc., San Diego, CA) at the dilution of 1:20,000, followed by ExtrAvidin alkaline phosphatase conjugate and alkaline phosphatase substrate: p-nitro blue tetrazolium chloride enhanced 5-bromo-4chloro-3-indolyl phosphate (Bio-Rad). The membranes with the transferred rgp120 were incubated with the HRP-conjugated V5 antibody (Invitrogen) at the dilution of 1:7000, developed with SuperSignal West Pico reagents (Pierce) and the bands were visualized by exposure on X-ray film (Kodak, Rochester, NY). The band intensity was determined by digital imager system and analyzed by Image J software.

### Treatment of virus stock or rgp120 with glycosidases

Two different glycosidases were used for specific removal of N-glycans: peptide N-glycosidase F (PNGase F) and endo N-glycosidase H (Endo H) (Prozyme, Hayward, CA). PNGase F removes all N-glycans from gp120/41 glycoprotein whereas Endo H removes high-mannose and hybrid glycans. Twenty-fold concentrated virus stocks or rgp120 were pre-denatured for 10 min at 95-100°C in a denaturing buffer provided with each glycosidase. After samples were chilled on ice for 5 min, enzymes were added to the reaction (for PNGase F-treated samples, additional detergent was added). Samples were incubated at 37°C for 30 h, and then electrophoretically separated under reducing conditions by SDS-PAGE and further analyzed by western blotting.

### Glycomic profiling by high-resolution liquid chromatography mass spectrometry (LC-MS)

Equal amounts of each rgp120 preparation (10 µg) were separated on a 4-15% Mini-PROTEAN TGX precast gel (Bio-Rad Laboratory Inc.) under denaturing and reducing conditions. Protein bands (∼130 kDa) were cut and digested with different proteases and glycosidases. Approximately 250 ng of peptides were loaded onto a 11-cm, 100 μm-diameter pulled tip packed with Jupiter 5 μm C18 reversed phase beads (Phenomenex, Torrance, CA). Samples were separated at a flow rate of 650 nL/min over 90 min on an Eksigent MicroAS autosampler and 2D LC nanopump (Eksigent, Dublin, CA). Mobile phases consisted of solvent A: 2.5% acetonitrile (ACN) and 0.1% formic acid (FA) and solvent B: 97.5% ACN and 0.1% FA. The following multistep gradient was used: 5 min, 2-5% B; 50 min, 5-35% B; 10 min, 35-40% B; 10 min, 40-60% B; 5 min, 60-98% B and held at 98% B for 10 min before re-equilibration. The eluted peptides were electrosprayed at 2 kV into a dual linear quadrupole ion trap Orbitrap Velos Pro mass spectrometer (MS) (Thermo Fisher, San Jose, CA). The MS was set to switch between a full scan (400 < m/z > 2000) followed by successive CID MS/MS (200 < m/z > 2000) scans of the most abundant precursor ions. A dynamic exclusion setting was set to exclude ions for 2 min after a repeat count of three within a 45-s duration.

All MS and MS/MS data was processed using the Byonic (Protein Metrics) and Pinnacle (Optys) software for N-glycopeptide site confirmation and to obtain extracted ion chromatograms (XIC) for every glycopeptide. The resultant XIC for N-glycopeptides >1E4 were used to calculate area under the curve for each glycopeptide ion. For each NGS the observed N-glycoforms were expressed as a percent relative abundance of the total sum of all glycoforms. Each NGS was classified as having predominantly high mannose, predominantly processed or mixed N-glycan heterogeneity profiles (Hargett *et al*, 2019). To determine the extent of processed glycans, a ratio of the area under the curve of the XIC for Man_9_ to Man_5_ was compared for predominantly high-mannose glycans, and a ratio of the XIC of HexNAc_4_Hex_3_Fuc_1_ to HexNAc_4_Hex_5_Fuc_1_ was compared for predominantly processed glycans.

### Molecular Dynamics Simulation

All four molecular dynamics simulations were carried out to characterize the fully glycosylated WEAU Env trimer variants under physiological conditions. The X-ray structure 5FYK (Stewart-Jones *et al*, 2016) was used as an initial atomistic model to model the four different WEAU Env trimer sequences on. Missing loops were built using loopy (Xiang *et al*, 2002). N-linked Man_5_ were modeled with *Glycosylator*, an in-house developed software. The program first identified the glycan species that were crystallographically resolved at each sequon. Mannose moieties were then added or removed from these glycans to create a Man_5_. Afterwards, the trimers were solvated in a 15Å padding water box and neutralized by the addition of NaCl at a concentration of 150 mM. The final systems were composed of about half a million atoms. The molecular dynamics simulations were performed using NAMD2.12 engine (Phillips *et al*, 2005), with the CHARMM36 force field(Best *et al*, 2012). TIP3P water parameterization was utilized to describe the water molecules (Jorgensen & Madura, 1983). The periodic electrostatic interactions were computed using particle-mesh Ewald (PME) summation with a grid spacing smaller than 1 Å. Constant temperature was imposed by using Langevin dynamics with a damping coefficient of 1.0 ps. Constant pressure of 1 atm was maintained with Langevin piston dynamics, 200 fs decay period and 50 fs time constant. Minimization and equilibration steps were performed as previously reported (Lemmin *et al*, 2017). Unrestrained molecular dynamics were performed up to 500 nanoseconds. The network analyses were carried out using the NetworkX library (Hagberg, 2008). Each glycan corresponded to one node in the graph. It has previously been shown that a correlation exists between the number of crystallographically defined glycan units and the number of sequons within a 50 Å sphere (Stewart-Jones *et al*, 2016). Therefore, any nodes, which sequons were within 50 Å of each other, were connected in the graph. The average non-bonded energy (van der Waals and electrostatic) was measured with VMD software (Humphrey *et al*, 1996). The edges between glycans that interacted with less than 0.1 kcal/mol were removed. Finally, each edge was weighted with the inverse of the interaction energy.

### Statistics

The infectivity of WEAU Env-pseudotyped viruses was compared by using two-tailed Student t test. Differences were considered to be significant at *P* values of less than 0.05.

## Supporting information

Appending Sup figs 1-3

WEAU-d16 MDS

WEAU-d16-N264S MDS

WEAU-d391 MDS

WEAU-d391-S264N

## Data Availability

The mass spectrometry proteomics data have been deposited to the ProteomeXchange Consortium via the PRIDE [1; http://www.ebi.ac.uk/pride] partner repository with the dataset identifier PXD017941.

[1] Perez-Riverol Y, Csordas A, Bai J, Bernal-Llinares M, Hewapathirana S, Kundu DJ, Inuganti A, Griss J, Mayer G, Eisenacher M, Pérez E, Uszkoreit J, Pfeuffer J, Sachsenberg T, Yilmaz S, Tiwary S, Cox J, Audain E, Walzer M, Jarnuczak AF, Ternent T, Brazma A, Vizcaíno JA (2019). The PRIDE database and related tools and resources in 2019: improving support for quantification data. Nucleic Acids Res 47(D1):D442-D450 (PubMed ID: 30395289).

### Acknowledgements

Support for this work was provided by the National Institutes of Health (NIH) grants GM098539, by Intramural Research Program of the Vaccine Research Center, NIAID, NIH, and by NIH T32 fellowship GM008111. This work was also supported in part by the University of Alabama at Birmingham (UAB) Center for AIDS Research (AI027767) developmental grant and a pilot grant from UAB School of Medicine. MR was supported in part by the UP grant IGA_LF_2020_016 and a grant of Ministry of School and Education, Youth and Sport of the Czech Republic, CZ.02.1.01/0.0/0.0/16_025/0007397. BK was supported with Federal funds from the National Cancer Institute, National Institutes of Health, under Contract No. HHSN261200800001E. The content of this publication does not necessarily reflect the views or policies of the Department of Health and Human Services, nor does mention of trade names, commercial products, or organizations imply endorsement by the U.S. Government. The authors would like to thank Beatrice Hahn, George Shaw, and Jiri Mestecky for helpful discussions.

## Author Contributions

Z.M., M.R., M.B.R., J.N. designed the study. B.F.K., M.R., J.N. selected WEAU gp120 variants from early and late stages of HIV-1 infection. Q.W., A.A.H., B.K., A.D., S.H., R.B. carried out the research. S.L.H., M.S.S. provided materials. R.R. prepared, carried out, and analyzed the MDS. C.H.S performed network analysis of MDS trajectories. S.K.F. assisted in analyzing glycan flexibility in MDS trajectories. O.K., G.Y.C., P.D.K., Z.M., M.R., M.B.R., J.N. supervised the research. Q.W., A.A.H., Z.M., M.R., M.B.R., J.N. drafted the manuscript. Q.W., A.A.H., B.K., R.R., B.F.K., O.K., P.D.K., Z.M., M.R., M.B.R., J.N. edited the manuscript. All authors analyzed data and commented on the drafts.

## Expanded View Figure Legends

**Figure EV1.**
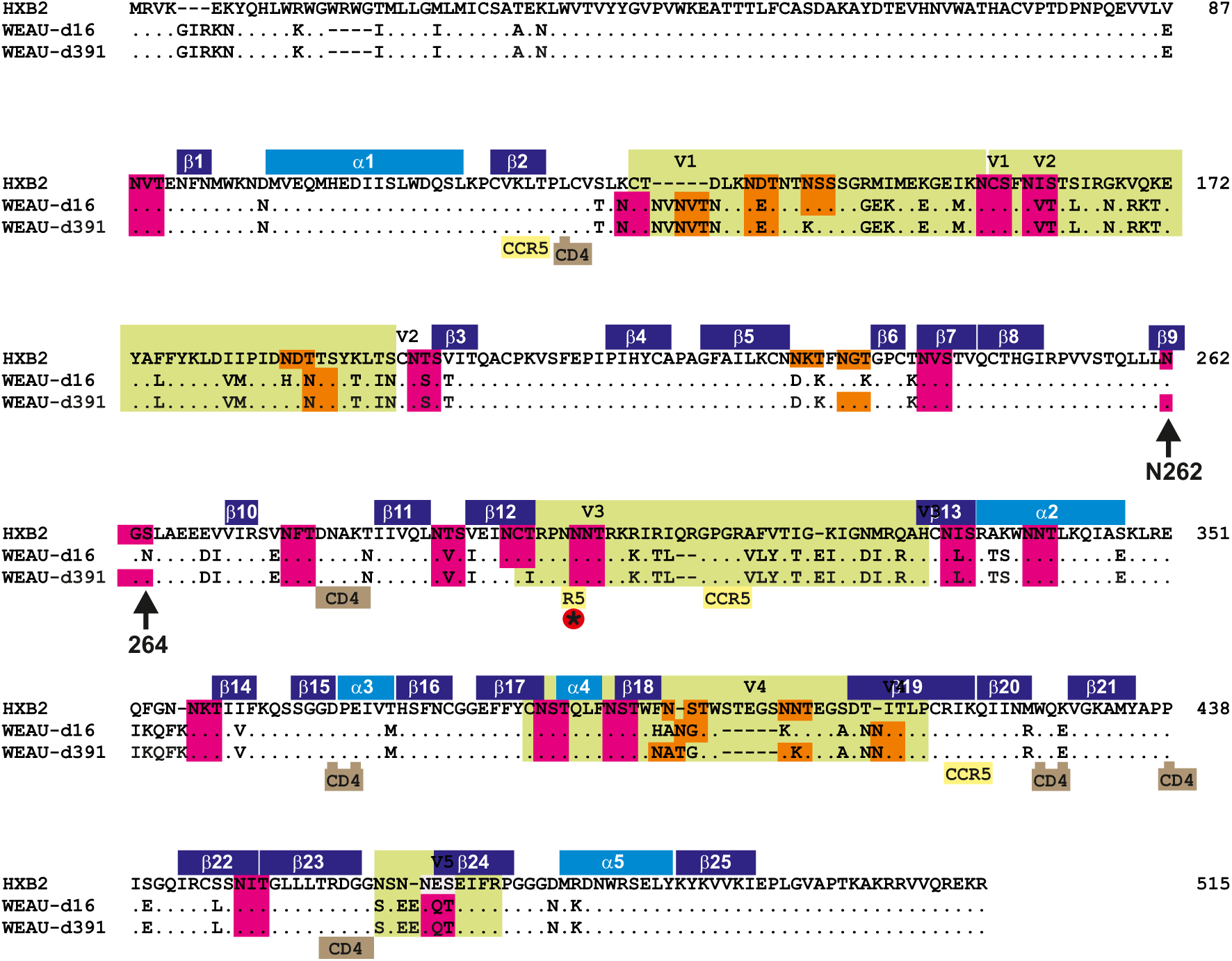
Alignment of gp120 segments of gp160 sequences and potential N-glycosylation sites (NGS) of HXB2 and WEAU-d16 and WEAU-d391. Secondary structures, α1 – α5 helices (light blue) and β1 – β25 strands (dark blue), are positioned according to published crystal structure of HIV-1_HXB2_. Variable loops V1 – V5 are in light-green color. Potential NGS are in pink, and shifting potential NGS are in orange color. CD4- and CCR5-binding regions are in brown and yellow, respectively. * indicates potential NGS 301 site associated with R5-to X4 switch.

**Figure EV2.**
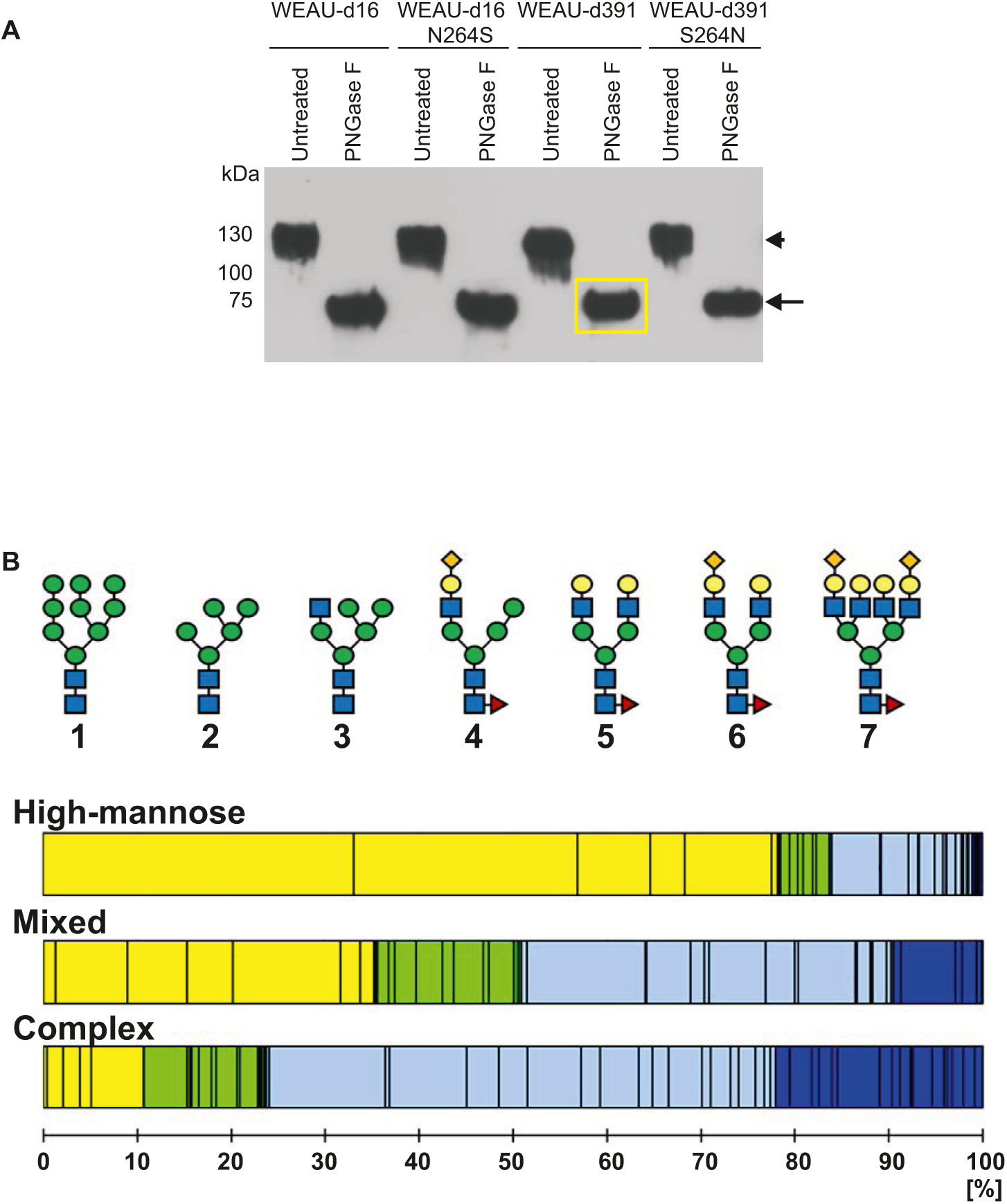
Normalization of rgp120 samples for glycomic analyses by mass spectrometry and examples of glycosylation heterogeneity. A Samples (untreated or PNGase F-treated) were analyzed by SDS-PAGE under denaturing and reducing conditions, transferred onto a PVDF membrane, and the gp120 detected by an anti-V5 tag antibody. The band intensity of PNGase F-treated samples was measured by digital imager system and analyzed by Image J software. One sample was selected as the standard control (in yellow box) and the amounts of other proteins were then adjusted accordingly. Arrowhead marks position of gp120 (∼120 KDa), arrow marks position of gp120 after deglycosylation with PNGase F (∼60 KDa). B Examples of the three types of the site-specific quantitative profiles observed in Env glycoproteins^27^: predominantly high-mannose, mixed, and predominantly complex (i.e., processed), shown as weighted distribution of high-mannose (yellow), hybrid (green), and complex (blue) oligosaccharides. Darker shading represents hybrid and complex N-glycans that contain sialic acid residue(s). Examples of the corresponding glycan structures are shown above the bars: 1,2; high-mannose glycans; 3,4; hybrid glycans; and 5,6,7; complex glycans. 1, Man_9_ high-mannose glycan; 2, Man_5_ high-mannose glycan; 3, hybrid glycan with single GlcNAc on Man residue; 4, extended version of glycan # 3 with added galactose and sialic acid and core fucose; 5, biantennary complex glycan with two galactose residues and core fucose; 6, biantennary complex glycan with two galactose residues, one sialic acid residue, and core fucose; 7, tetraantennary glycan with four galactose residues, one sialic acid residue, and core fucose. Symbols for monosaccharides: blue square, GlcNAc; green circle, mannose; red triangle, fucose; yellow circle, galactose; orange diamond, sialic acid.

**Figure EV3.**
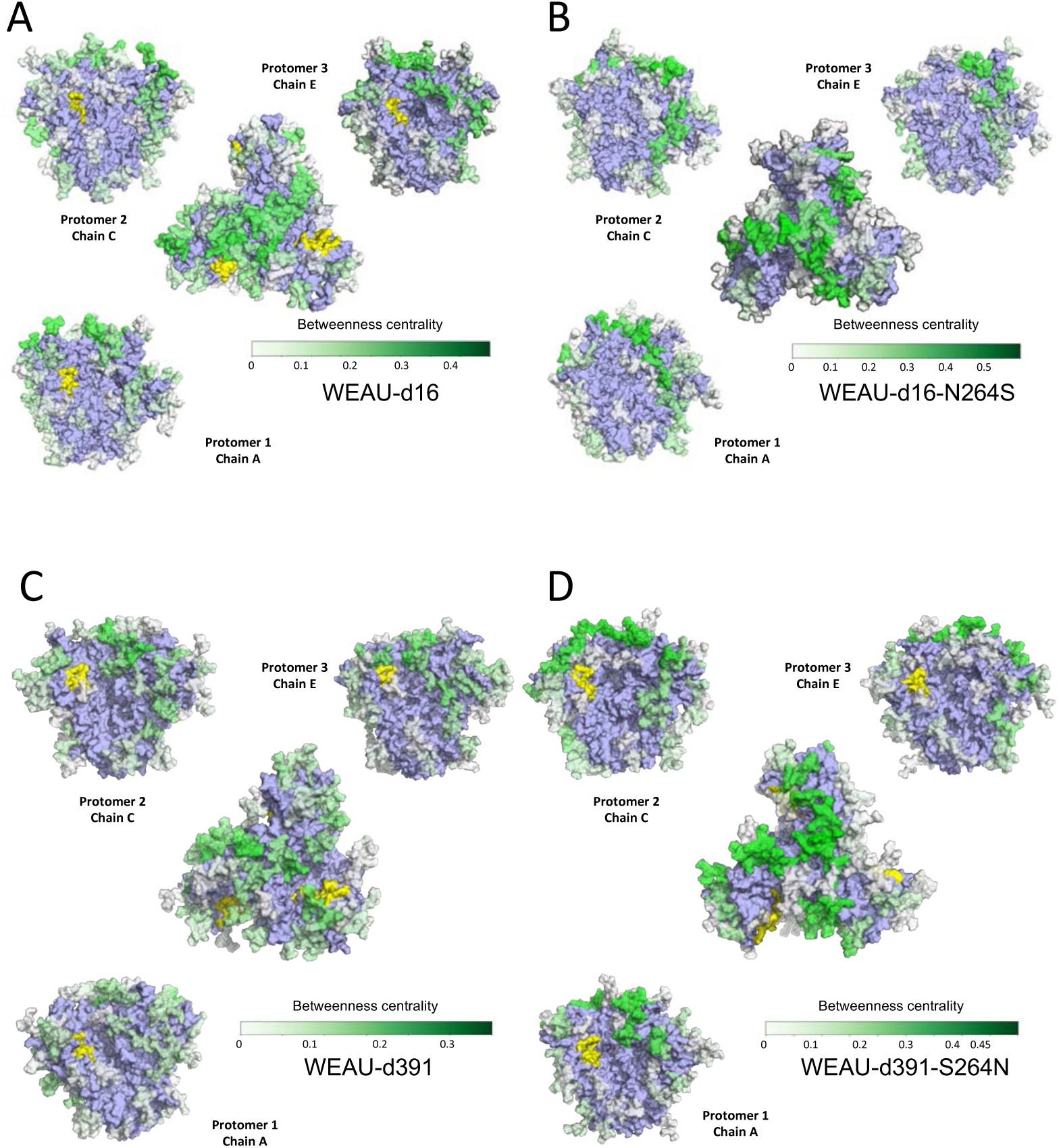
WEAU gp120 trimers betweenness centrality heatmaps. A Betweenness centrality value of each oligosaccharide from WEAU-d16 Env mapped onto the structural model. The CD4 binding site is highlighted in yellow and the Env is colored in light blue. Each glycan is colored based on its betweenness centrality value. B Betweenness centrality value of each oligosaccharide from WEAU-d16-N264 Env mapped onto the structural model. The CD4 binding site is highlighted in yellow and the Env is colored in light blue. Each glycan is colored based on its betweenness centrality value. C Betweenness centrality value of each oligosaccharide from WEAU-d391 Env mapped onto the structural model. The CD4 binding site is highlighted in yellow and the Env is colored in light blue. Each glycan is colored based on its betweenness centrality value. D Betweenness centrality value of each oligosaccharide from WEAU-d391-S264N Env mapped onto the structural model. The CD4 binding site is highlighted in yellow and the Env is colored in light blue. Each glycan is colored based on its betweenness centrality value.

**Figure EV4.**
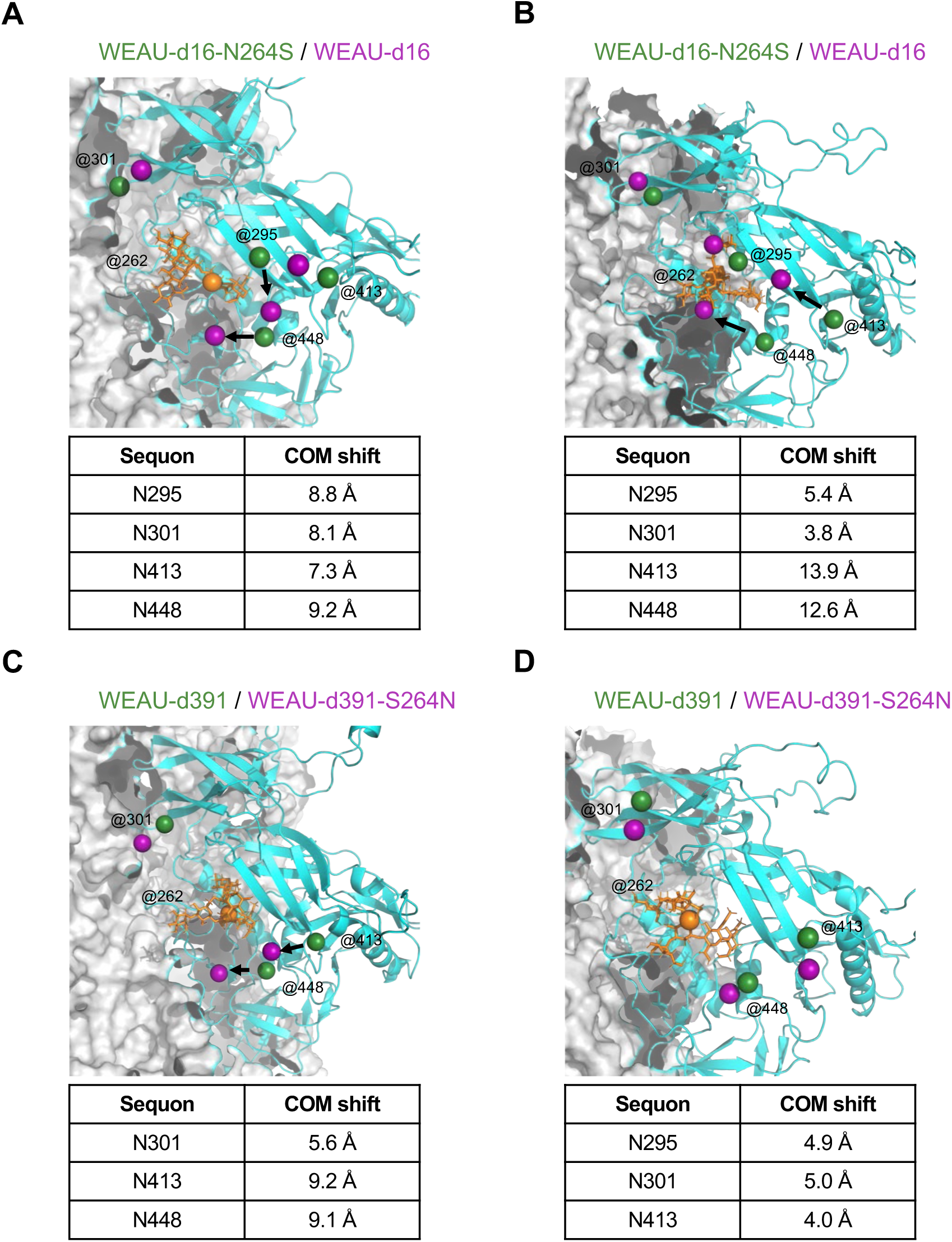
Center of Mass (COM) for glycans within the high-mannose patch highlight glycan movement when glycan 262 is absent. A COM differences were assessed for each oligosaccharide between WEAU-d16-N264S (purple) and WEAU-d16 (dark green) for protomer 1. B COM differences were assessed for each oligosaccharide between WEAU-d16-N264S (purple) and WEAU-d16 (dark green) for protomer 3. C COM differences were assessed for each oligosaccharide between WEAU-d391 (purple) and WEAU-d391 S264N (dark green) for protomer 1. D COM differences were assessed for each oligosaccharide between WEAU-d391 (purple) and WEAU-d391 S264N (dark green) for protomer 3.

