## Appending Sup figs 1-3 for "Impact of glycan positioning on HIV-1 Env glycan shield density, function, and antibody recognition"

**Appendix Legends**

**Appendix Fig S1. Infection of TZM-bl reporter cells with WEAU-d16 and WEAU-d391 Env-pseudotyped viruses produced by 293F or 293T cell lines.**

A Using plasmids encoding *gp160* genes from WEAU-d16 and WEAU-d391 and *env*-deficient HIV-1 backbone vector (pSG3$_{\Delta Env}$), we transfected HEK293 cells and produced Env-pseudotyped viruses, WEAU-d16 and WEAU-d391. For virus production, we compared the conventional adherent 293T cells grown in serum-supplemented medium and FreeStyle 293 (293F) cells grown in suspension in serum-free medium. Infectivity, calculated as relative light units (RLU) normalized to viral load (p24), was similar for each construct irrespective of the producing cell line.

B Amount of p24 in the viral stocks. The data, from three and four independent experiments, respectively, are shown as mean ± standard deviations. We found comparable amounts of p24 in all produced virions. Based on these results, Env-pseudotyped viruses used in this study were produced in 293F cells.

**Appendix Fig S2**. **Identification of gp160 and gp120 from WEAU-d391 virions**.

Western blots of 20-times concentrated viral stocks separated on SDS-PAGE, before and after treatment with PNGase F, were probed with:

A IgG isolated from serum of an HIV-1-infected individual;

B gp41-specific monoclonal antibody 2F5 (reacts with gp160 but not gp120).

Red and blue arrows mark gp160 and gp120, respectively. After digestion with PNGase F, bands corresponding to the heavily glycosylated gp160 and gp120 disappeared and new bands, corresponding to the deglycosylated gp160 (~80 kDa) and gp120 (~60 kDa) polypeptides were detected. Analysis of WEAU-d16 virions provided very similar results (data not shown), confirming that all virions were similar in their gp120 and gp160 contents.

**Appendix Fig S3**. **Neutralization susceptibility of WEAU-d16 and WEAU-d391.**

A. Neutralization of Env-pseudotyped viruses by IgG isolated from plasma of WEAU subject 5 years after HIV infection.

B Neutralization of Env-pseudotyped viruses by monoclonal BnAb 2G12.

**Supplementary movies:**

We are showing only one gp120 chain (chain C) per trimer of WEAU-d16, WEAU-d16-N264S, WEAU-d391, WEAU-d391-S264N Env variants, but the behavior is quite similar for all chains, as the COM analysis showed. Man5 is modeled at the NGS shown. The movies, based on unrestrained molecular dynamics performed for up to 500 nanoseconds as detailed in Supplemental Methods, are focused on the HMP glycans.

Color code for glycans is as follows:

Orange: N262 (if present)

Blue: N295 (if present)

Red: N301

Yellow: N332

Green: N413

Pink: N448

### Appendix Fig S1

**A**

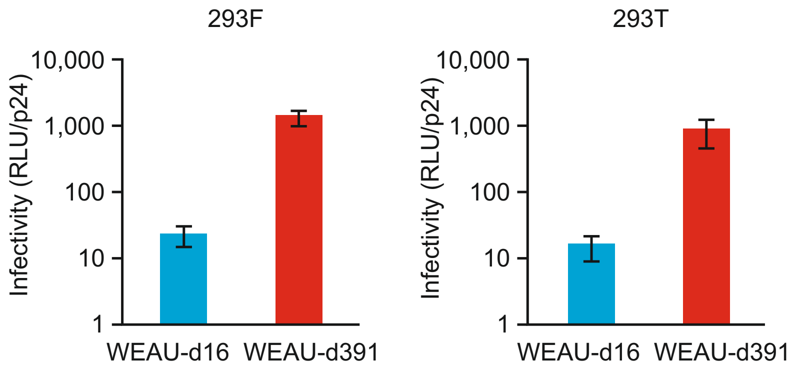


**B**

HIV-1 p24 in viral stocks

| | **293F[a] (pg/ml)** | **293T (pg/ml)** |
|---|---|---|
| WEAU-d16 | 252 (n=4) | 206 |
| WEAU-d391 | 276 (n=3) | 209 |

[a] mean values from individual stocks

### Appendix Fig S2

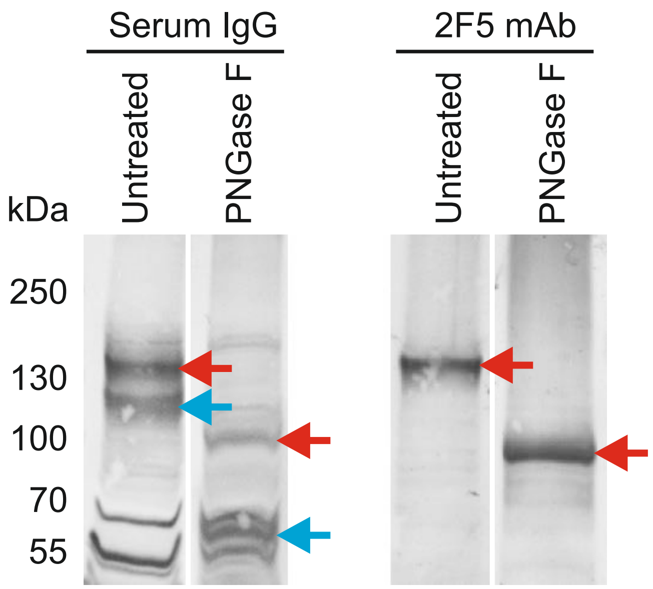

### Appendix Fig S3

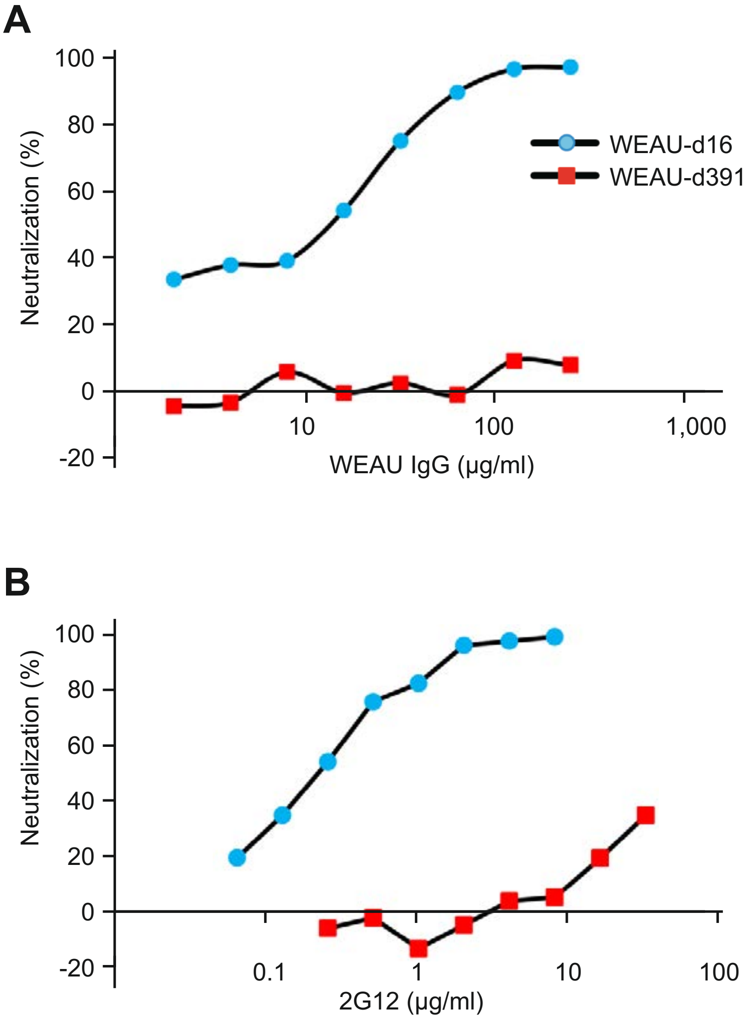
